# Large and interacting effects of temperature and nutrient addition on stratified microbial ecosystems in a small, replicated, and liquid dominated Winogradsky column approach

**DOI:** 10.1101/2020.12.09.415786

**Authors:** Marcel Suleiman, Yves Choffat, Uriah Daugaard, Owen L Petchey

**Author notes:** Corresponding author: Marcel Suleiman, Department of Evolutionary Biology and Environmental Studies, University of Zurich, Switzerland.

## Abstract

Aquatic ecosystems are often stratified, with cyanobacteria in oxic layers and phototrophic sulfur bacteria in anoxic ones. Changes in stratification caused by global environmental change are an ongoing concern. Increasing understanding how such aerobic and anaerobic microbial communities, and associated abiotic conditions, respond to multifarious environmental changes is an important endeavor in microbial ecology. Insights can come from observational and experimental studies of naturally occurring stratified aquatic ecosystems, from theoretical models of ecological processes, and from experimental studies of replicated microbial communities in the laboratory. Here we demonstrate a laboratory-based approach with small, replicated, and liquid dominated Winogradsky columns, with distinct oxic/anoxic strata in a highly replicable manner. Our objective is to apply simultaneous global change scenarios (temperature, nutrient addition) on this micro-ecosystem to report how the microbial communities (full-length 16SrRNA-seq.) and the abiotic conditions (O_2_, H_2_S, TOC) of the oxic/anoxic layer responded to these environmental changes. Composition of the strongly stratified microbial communities was greatly affected by temperature and by the interaction of temperature and nutrient addition, demonstrating the need of investigating global change treatments simultaneously. Especially phototrophic sulfur bacteria dominated the water column at higher temperatures, and may indicate the presence of alternative stable states. We show that the establishment of such a micro-ecosystem has potential to test global change scenarios in stratified eutrophic limnic systems.

## Introduction

Microorganisms are key players in nearly all ecosystems, ranging from the human gut to marine and freshwater habitats. The functioning of microbial communities is critical for many ecosystem services, since microorganisms are the driving force behind biogeochemical cycles (Kertesz 2000; Weiss et al. 2003; Falkowski et al. 2008; Stein and Klotz 2016). Nevertheless, microbes are highly dependent on environmental conditions; even slight changes can lead to taxonomic and functional community shifts (Allison et al. 2013; Evans and Wallenstein 2014). Consequently, microbial communities and their functions are significantly affected by global change (Cavicchioli et al. 2019), which has recently led to the emergence of the research field termed “Global Change Microbiology” (Boetius 2019).

Stratification of productive lakes is a common phenomenon observed in summer, resulting in warm and oxygen-rich upper water layers, dominated by cyanobacteria and algae, and colder anoxic deeper layers, that harbor heterotrophic biomass-degraders and sulfate-reducing bacteria. In between these layers, the metalimnion can be found, which is formed by decreasing light-intensities and oxygen concentrations (microaerophilic), and an increasing pool of reduced sulfur compounds, creating ideal niches for phototrophic sulfur bacteria and chemolithoautotrophic microorganisms (Jorgensen et al. 1979; Morrison et al. 2017; Savvichev et al. 2018; Vigneron et al. 2021).

Beside chemical and physical parameters that influence the presence of oxic and anoxic layers, it was recently postulated that mutual inhibition between cyanobacteria and anaerobic sulfur-dependent bacteria creates and maintains the distinct oxic and anoxic zones of such ecosystems (Bush et al. 2017). The mathematical model used in that study predicts abrupt transitions between aerobic and anaerobic microorganisms and the occurrence of oxic-anoxic regime shifts. Moreover, studies have demonstrated that global change consequences could have strong impacts on this sensitive ecosystem, including dominant growth of harmful cyanobacteria due to warming (Posch et al. 2012), or increasing primary production and creation of anoxic layers due to increased nutrient input (Luek et al. 2017).

The mentioned studies show that important insights come from analyses of observations of naturally occurring aquatic ecosystems (Vigneron et al. 2021), and from theoretical models of the plausible and relevant ecological and biochemical processes. In addition, in the field of global change biology, targeted experiments that employ a limited number of standardised synthetic model ecosystems have been described as an “entirely different approach” (Hutchins et al. 2019) and are considered vital alongside studies of naturally occurring ecosystems (Lahti et al. 2014) and hybrids of both approaches (De Vos et al. 2017). Key challenges in global change microbiology that are amenable to research with synthetic ecosystems include the role of evolutionary processes, the role of historical contingency (Widder et al. 2016) and interactions among microorganisms (Overmann and van Gemerden 2000), and the integration of data and theory (Widder et al. 2016). Experiments about the role of environmental, organismal interactions and biochemical feedbacks in stratified aquatic micro-ecosystems appear absent, however.

Establishing suitable such standardised-model ecosystem for analyzing potential responses of a broad range of microorganisms remains crucial. Being able to monitor global change responses of diverse functional groups of potentially interacting microbes is a desirable feature of such an experimental system, which point towards systems with strong abiotic gradients in space and or time. In this work, we get inspiration from an “old” but highly valuable approach, the Winogradsky column (Zavarzin 2006; Dworkin 2012; Pagaling et al. 2017). We constructed a modified and smaller version that allows a highly replicable development of a broad range of complex and dynamic microbial groups in one experimental unit. In contrast to classical Winogradsky columns, our micro-ecosystems are mostly liquid, with only a small sediment layer (∼ 6 % v/v of the column). Thus, we created a highly replicable liquid oxic-anoxic interphase and self-developing model systems.

By applying this approach, we were able to analyze microbial responses to multifarious environmental change (temperature and nutrient addition manipulated factorially), including the responses of community composition and abiotic environmental conditions. As well as showing how microbial community composition, dissolved oxygen, pH, and hydrogen sulphide respond to temperature and nutrient addition, we highlight the potential for global change microbiology research of this new approach coupled with state-of-the-art sequencing technologies.

We hypothesize that higher temperature will increase the dominance of anaerobic microbes due to an increase in the extent of the anoxic zone, due to the lower solubility of oxygen in warmer water. We hypothesize that nutrient addition (addition of ammonium phosphate, highly used fertilizer (Cao et al. 2018)) will alter community composition, but we cannot *a priori* say how, as this would depend on features of the organisms, such as competitive and facilitative interactions among them, which we do not have sufficient information about. We did not have any *a priori* expectation of whether the combined effect of temperature and nutrient addition would be additive (no interaction), more than additive (positive interaction), or less than additive (negative interaction). We anticipated the potential for micro-ecosystems to become entirely oxic or entirely anoxic, and for changes in composition to be non-linear and to potentially include discontinuous responses that are predicted by theory.

## Materials and methods

### Preparation of micro-ecosystems

The vessels used to house the micro-ecosystems were standard glass test-tubes (diameter 13 mm, height 16 cm). Each was equipped with two oxygen sensors (PreSens Precision Sensing GmbH, Germany), one at a height of 4 cm (bottom sensor) and the other at 14 cm (top sensor), both stuck to the inside surface of the test-tube wall. These sensors allow optical measurements of oxygen to be taken through the wall of test tube.

Each test-tube (which we also refer to as a “column”) had a butyl rubber stopper on top, with two canulas. The longer canula (StericanR, length 120mm, diameter 0.8 mm) reached to the depth of the bottom O_2_ sensor and was covered with an oxygen-protecting plug (BD Plug). The shorter canula (length 40 mm, 0.9 mm diameter) was always open and extended only into the headspace of the micro-ecosystem. This shorter canula therefore allowed for gas exchange between the headspace and the atmosphere (Fig. 1a).

**Fig. 1.**
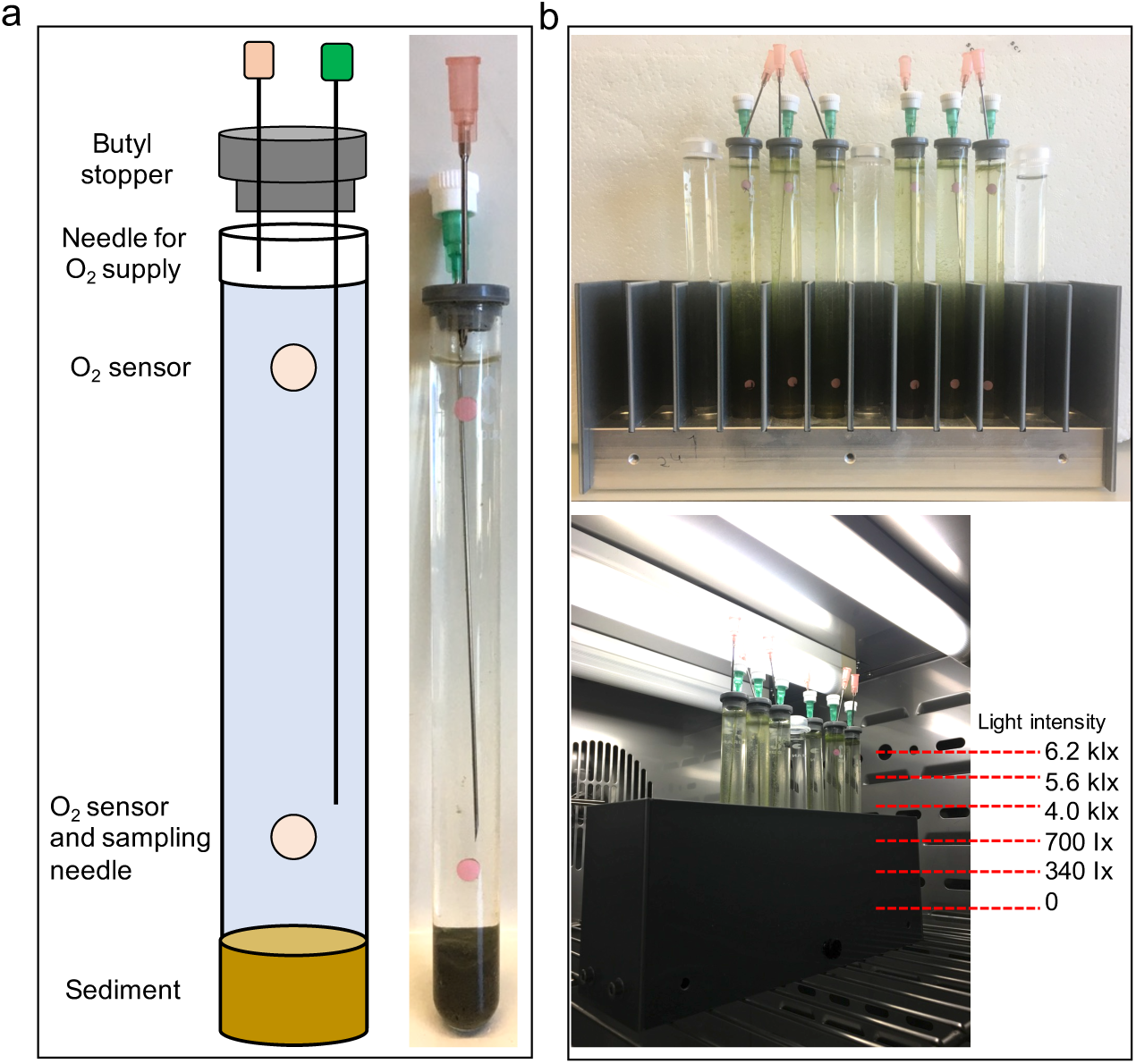
Preparation and incubation conditions of the micro-ecosystems. (**a**) Columns were filled with 1.5 cm sediment and pond water with supplementary resources (see main text for details). Columns were equipped with two oxygen sensors (top and bottom), and closed with a butyl rubber stopper. One canula allow gas exchange between the headspace and the atmosphere (pink). A second canula is used for taking samples at the bottom sensor (green, anoxic stopper). (**b**) Micro-ecosystems were incubated in light-gradient-producing holders. Sediment was covered inside the metal base (1.5 cm depth) for incubation without light. Columns were thereby incubated with a light gradient ranging from 6.2 kilo Iux (klx) to 0 Iux (lx).

Micro-ecosystems were established from sediment and water samples taken in July from a small pond in Zürich, Switzerland (47°23′51.2″N 8°32′33.3″E, temperature of 24.6 °C, pH of 6.7.) at a depth of 30 cm. In total, 200 g sediment and 2 L water were collected. The sediment sample was supplemented with sterile 0.5 % crystalline cellulose, 0.5 % methyl-cellulose, 1 % CaSO_4_, 0.2 % CaCO_3_ and 0.01 % NH_4_H_2_PO_4_. Sediment was homogenized by mixing for 30 min, before each test-tube was filled with sediment up to a height of 1.5 cm (in 0.5 cm steps, with constant mixing of the source sediment bottle between steps). Sediment that stuck to inner wall of the test-tube was removed carefully in all cases. Columns were filled with well-homogenized 16 mL pond water (supplemented with 0.01 % NH_4_H_2_PO_4_), leaving approximately 1 cm of headspace. In contrast to classical Winogradsky columns (Zavarzin 2006; Dworkin 2012; Rundell et al. 2014), our set-up is dominated by liquid and contains a small amount of sediment. Columns were incubated at room temperature for two hours without light to allow the sediment to settle fully, after which an initial oxygen concentration of the top and bottom sensor was recorded and the columns were placed in incubators.

### Incubation and treatments

Each column was incubated at one of seven temperatures: 12, 16, 20, 24, 28, 32 or 36 °C. Incubators (Pol Eco Apparatura sp.j.) were equipped with two fluorescent light sources (PHILIPS Master TL8W/840) with a dark-light cycle of 8:16 hours. Columns were placed in a 1.5 cm hole in a metal base, so that the sediment layer was continuously kept in darkness. Up to a height of 10 cm, the columns were surrounded by a plastic holder that causes a gradient of light, ranging from 340 Ix at the intersphere of sediment:water at the bottom and up to 700 Ix at the top of the light-protector (Fig. 1b). The top of the column experienced fluorescent light intensity of 6200 Ix. Each holder contained six columns, in two groups of three replicates, separated by a water-filled column. This arrangement was placed in the middle of both light sources in the incubator, allowing the same light-intensity at the back and front of each column.

A second treatment was the manipulation of nutrient concentration. For this, 0.1 % NH_4_H_2_PO_4_ was added to half of the columns by taking 16 µL of a sterile 10 % NH_4_H_2_PO_4_ solution, while 16 µL sterile water was used for the controls. This treatment was performed once per week (day 1, 8, 14, 20) and was factorially crossed with the temperature treatment, resulting in 7 (temperature) x 2 (nutrient levels) and 3 replicated, giving a total of 42 columns. These were incubated for 22 days.

### Sampling and measurements

Liquid samples were taken on day 8, 14 and 20 by collecting 200 ul of liquid from the bottom of the columns by using the installed canula.This liquid was used to measure H_2_S (Microsensor, AMT Analysenmesstechnik GmbH, Germany) and was replaced with anoxic sterile lake water. Non-invasive oxygen measurements were performed at the top and bottom sensor of the columns (everyday except of day 5, 11, 12 and 17). Oxygen concentration is presented in % saturation adjusted for the prevailing temperature and pressure (e.g. 21% saturation occurs when water is in equilibrium with the surrounding atmosphere). Thus the known effect of temperature on oxygen solubility does not affect our analyses.

After incubation, all columns were destructively sampled. For each column, the whole water column (16 mL) and the sediment layer were processed separately. One replicate of each treatment combination had the water column separated at the height of the bottom oxygen layer, giving a sample of the upper and lower liquid layer as well as a sample for the sediment. Each sample was centrifuged (16,000 rpm, 3 min) and pellets were frozen at −80 °C until DNA extraction. Supernatant was used for pH measurements, as well as total nitrogen and total organic carbon analysis. While sampling, the H_2_S concentration at the sediment-water interphase was measured under anoxic conditions.

### DNA extraction, 16S rRNA amplification

DNA extractions were performed with ZymoBIOMICS DNA Microprep Kit (ZymoResearch), following the manufacturer’s instructions. 16S rRNA gene amplifications were performed with 27F forward primer (5’-AGRGTTYGATYMTGGCTCAG-3’) and 1592R reverse primer (5’-RGYTACCTTGTTACGACTT-3’), resulting in an amplification product of ∼1500 bp. Primers were equipped with 5’phosphate and a 5’buffer sequence of GCATC. Barcodes, PCR chemicals and conditions (27 cycles) were performed as suggested by Pacific Biosciences (PacBio, 2020). PCR products were visualized on an 1 % (w/v) agarose gel and were pooled with equal concentrations. Afterwards, PCR products were purified using AMPure®PB beads (PacBio), following manufacturer’s guidelines. Besides 16S rRNA amplification for PacBio long read sequencing, also DGGE analysis performed as described previously (Antranikian et al. 2017; Suleiman et al. 2019) in order to identify the white bacterial population.

### Library Preparation and Sequencing

Sequencing was performed at the Functional Genomic Center Zürich, Switzerland. The SMRT bell was produced using the SMRTbell Express Template Prep Kit 2.0 (PacBio). The input amplicon pool concentration was measured using a Qubit Fluorometer dsDNA High Sensitivity assay (Life Technologies). A Bioanalyzer 12Kb assay (Agilent) was used to assess the amplicon pool size distribution. 0.75μg of amplicon pool was DNA damage repaired and end-repaired using polishing enzymes. A ligation was performed to create the SMRT bell template, according to the manufacturer’s instructions. The SMRT bell library was quality inspected and quantified using a Bioanalyzer 12Kb assay (Agilent) and on a Qubit Fluorimeter (Life technologies) respectively. A ready to sequence SMRT bell-Polymerase Complex was created using the Sequel II Binding Kit 2.1 and Internal Control 1.0 (PacBio) according to the manufacturer instruction. The Pacific Biosciences Sequel II instrument was programmed to sequence the library on 1 Sequel II SMRT Cell 8M, taking 1 movie of 15 hours per cell, using the Sequel II Sequencing Kit 2.0 (PacBio). Sequencing data quality were checked, via the PacBio SMRT Link software, using the “run QC module”. Sequencing raw data are deposited at NCBI SRA under the BioSample accession numbers SAMN16774060 to SAMN16774152.

### Bioinformatics

Bioinformatics were performed using R and the package Dada2 (Callahan et al. 2016). In short, reads were filtered regarding primer sequences, length (1300-1600 bp), quality, error rates and chimeras. The outcome of the filter processes can be observed in Supplementary data for each sample. Finally, the constructed sequence table was aligned with the SILVA ribosomal RNA database (Quast et al. 2012), using version 138 (non-redundant dataset 99). Afterwards, a phyloseq formatted data object was constructed, containing the amplicon sequence variant (ASV) table, taxonomy table and sample metadata, using the package phyloseq (McMurdie and Holmes 2013), which was also used for microbial community analysis together with the vegan package (Oksanen et al. 2019). All sequences that were assigned to “chloroplast” were removed from the phyloseq file, since primers were chosen for bacterial 16S rRNA gene and bind incorrectly to chloroplast sequences.

Microbial community composition was analysed with non-metric multidimensional scaling (NMDS), using the metaMDS function from the vegan package. NMDS was used because a principal component analysis of the compositional variation exhibited a strong “arch-effect”, which can hinder the use of ordination scores in further analyses that we wanted to make. The metaMDS function was instructed to use Bray-Curtis distance, to transform the data to have 2-dimensions. The metaMDS function was instructed to try 200 different starting positions in order to assure convergence. NDMS was performed on relative abundances of sequences with a relative abundance of at least 1% in at least one sample. The 2-dimensional NDMS solution converged with a stress of 0.15. A 3-D solution converged with stress of 0.11. This suggests the that 2-D solution is a good representation of the compositional variation among the samples, and that the 3-D solution is not much better. Linear models with appropriate error structures were used to assess the weight of evidence supporting causal effects of treatments and treatment interactions on response variables. All analyses involving NDMS and variables derived from it were repeated with a different standardization (centred log ratios) and a different distance measure (Aitchison distance) (Gloor et al. 2017). These different methods did not affect the conclusion of strong and relatively linear effect of temperature on composition (NMDS1), and of a nonlinear and interacting effect of temperature and nutrient treatment on composition (NMDS2).

We intentionally avoid referring to effects as significant or not significant, and rather use the terms: no evidence p > 0.1, weak evidence 0.1 > p > 0.05, moderately strong evidence 0.05 > p > 0.01, strong evidence 0.01 > p > 0.001, very strong evidence p < 0.001.

## Results

### Macroscopic observations

All columns developed a stratified growth of colorful microorganisms within the incubation time, regardless of the temperature and nutrient addition treatment (Fig. 2a & b). Macro- and microscopical observation of morphology and color indicated that the micro-ecosystems likely consisted of a relatively large upper layer of cyanobacteria and algae and phototrophic sulfur bacteria in a smaller lower layer. Moreover, a white layer occurred in all columns during the first 10 days (Fig. 2c) at different positions depending on the incubation temperature (12 °C and 16 °C below the bottom-sensor, and at higher temperatures above the bottom-sensor). DGGE analysis identified this layer as a community of species of the microaerophilic genus *Magnetospirillum*, indicating that this layer is the oxic-anoxic interphase of the micro-ecosystems. In general, all micro-ecosystems developed an oxic and anoxic part (Fig. 5), and triplicates of each treatment combination predominantly showed comparable growth and development. Gas formation was observed in the sediment layers of the columns, indicating an active microbial community (Fig. 2d).

**Fig. 2.**
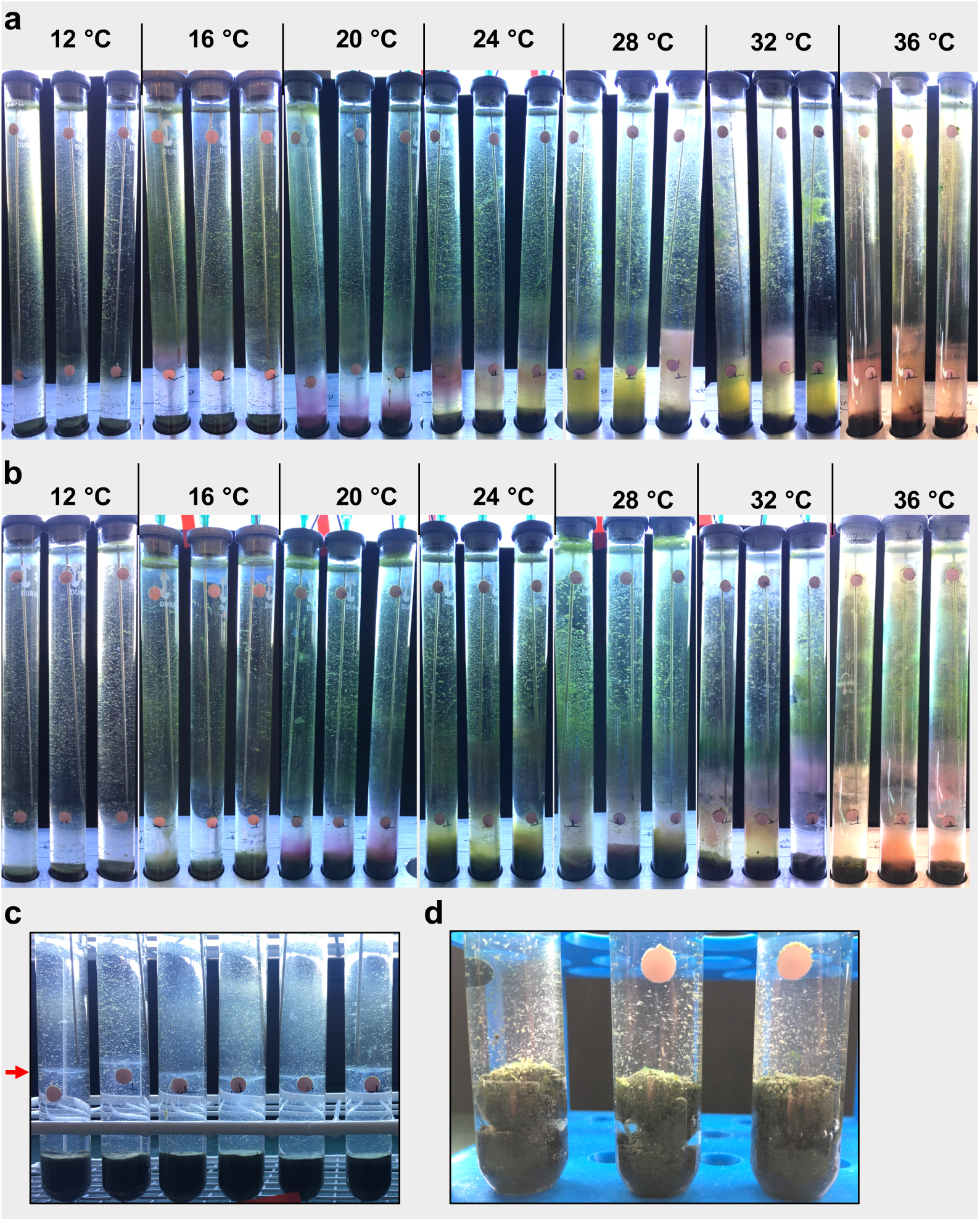
Macroscopical observation of the micro-ecosystems. **(a)** Control micro-ecosystems (triplicates) after incubation at 12-36°C for 22 days. **(b)** Nutrient addition micro-ecosystems (triplicates) after incubation at 12-36 °C for 22 days. **(c)** Observation of a white bacterial cloud within 10 days of incubation. Exemplary columns incubated at 20 °C of day 10 are shown. Red arrow highlights the position of cloud. **(d)** Gas formation in the sediments, exemplary picture taken at day 12 of columns incubated at 28 °C.

### Relative abundances of microbial taxa of the water column

In order to study the microbial communities in detail, full-length 16S rRNA sequencing was performed. Removal of sequences that did not occur at more than 1 % in at least one sample resulted in 343 different amplicon sequence variants (from initial 5633 unique sequences in all samples).

All of the micro-ecosystems at 12 °C were dominated by members of the order *Burkholderiales* (54 – 69 %) (Fig. 3a, Fig. S1 for details). Relative abundance of *Burkholderiales* decreased to 15 – 37 % (control) and 29 – 53 % (nutrient addition) at 16 °C, and relative abundance of *Rhodospirillales* increased up to 23 %. At 20 °C, members of *Chlorobiales* and *Chromatiales* became more abundant. Members of *Chlorobiales* were dominant in all micro-ecosystems at 24 °C (33 – 64 % controls, 62 – 64 % nutrient addition-treatments), 28 °C (42 – 92 % controls, 35 – 70 % nutrient addition-treatments), and 32 °C (78 – 90 % controls, 18 – 50 % nutrient addition). At 36 °C, members of *Chlorobiales* disappeared, and in no-nutrient-addition treatments, members of the cyanobacterial order *Limnotrichales* were dominant (up to 25 %; but none occurred in nutrient addition treatment).

**Fig. 3.**
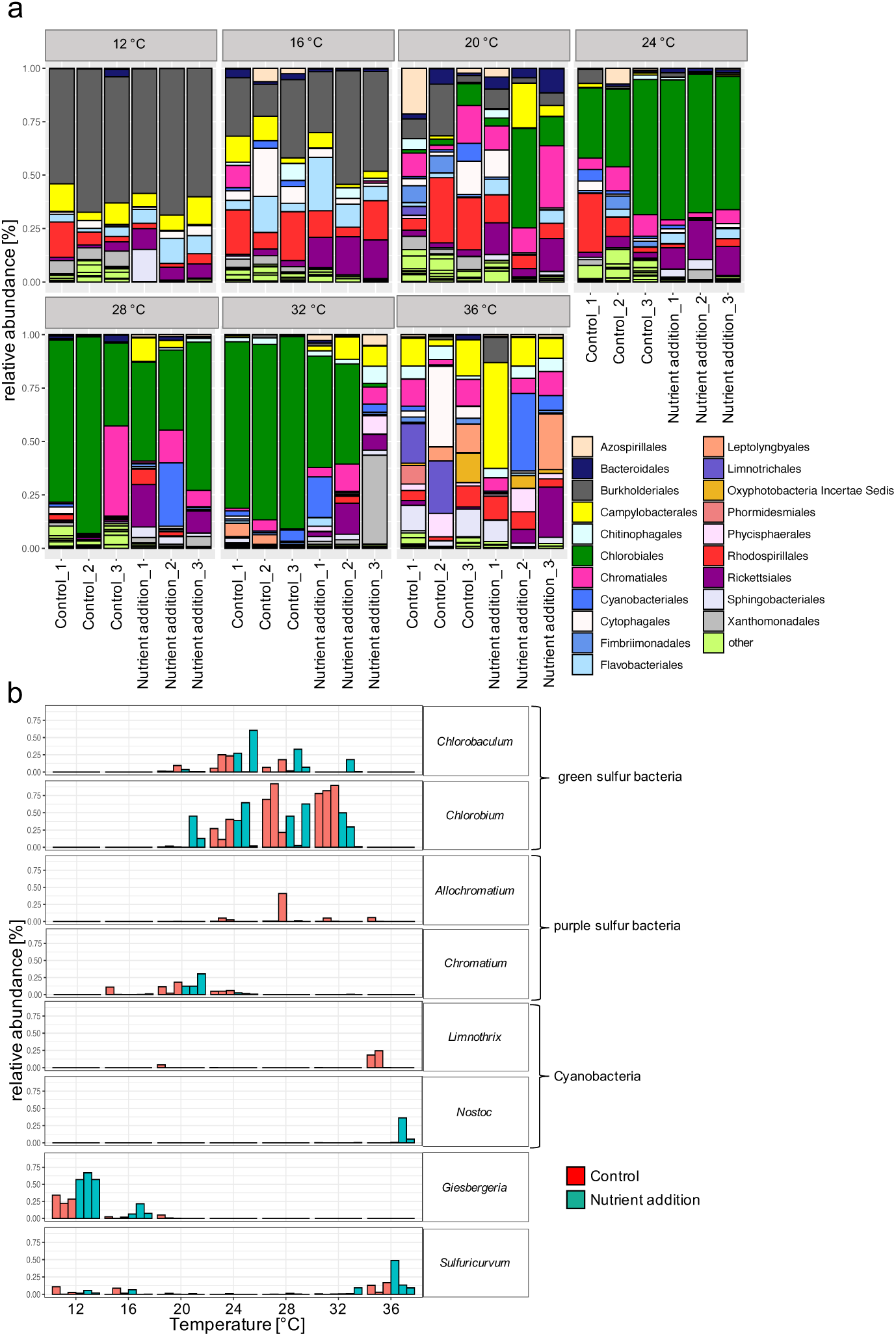
Microbial community composition of the micro-ecosystems. **(a)** Relative abundance on order level is shown for each micro-ecosystem. Orders with relative abundances < 7 % in all samples were assigned to “other”. **(b)** Relative abundance of 8 dominant and functional important genera depending on the temperature and treatment conditions. Red bars represent control triplicates, blue bars nutrient addition triplicates.

The full-length 16S rRNA amplicon sequencing resulted in a detailed account of the microbial community to genus level (Fig. 3b, Fig. S2-S8 for details). All micro-ecosystems at 12 °C (Fig. 3b, Fig. S2) were dominated by members of the aerobic genus *Giesbergeria* (controls 22 – 34 %, nutrient addition 57 – 67 %), followed by *Aquaspirillum* in the controls and *Flavobacterium* in the nutrient-addition micro-ecosystems. At 16 °C (Fig. 3b, Fig. S3), control micro-ecosystems were dominated by members of the aerobic genus *Uliginosibacterium* (12 – 27%), which had a very low relative abundance in the nutrient addition treatment (less than 2 %). In contrast, *Giesbergeria* and *Aquaspirillum* (4 – 35 %) were very abundant in the nutrient addition treatment.

Genus composition at 20 °C (Fig. 3b, Fig. S4) included anaerobic phototrophic sulfur bacteria, in particular the green sulfur bacteria *Chlorobium* appeared highly abundant in the nutrient addition treatments (up to 46 %) and the purple sulfur bacteria *Chromatium* in all micro-ecosystems at this temperature (up to 28 %). With increasing temperature, green sulfur bacteria *Chlorobium* and *Chlorobaculum* genera increased in relative abundance. At 24 °C (Fig. 3b, Fig. S5), *Chlorobium* and *Chlorobaculum* were detected in the controls, with 11 – 39 % and 6 – 25 % relative abundance, respectively, and in the nutrient addition treatments with 39 – 65 % and 27 – 60 %. Furthermore, the picocyanobacterial genus *Cyanobium* appeared (up to 7 %) in the control micro-ecosystems.

At 28 °C (Fig. 3b, Fig. S6), green sulfur bacteria continued to increase their dominance, especially *Chlorobium* (up to 92 %). Interestingly, the *Control_3* column consisted of a comparable lower amount of green sulfur bacteria, but additionally of the purple sulfur bacteria *Allochromatium* (41 %). This finding can be confirmed by the macroscopical observation of the pink layer (Fig. 3a). Additionally, the *Nutrient addition_2* column consisted of *Chlorobaculum* instead of *Chlorobium* in contrast to the *Nutrient addition_1* and *Nutrient addition_3* columns.

At 32 °C (Fig. 3b, Fig. S7), *Chlorobium* dominated the control-micro-ecosystems, while this genus also had lower though still high relative abundance in the nutrient addition treatments (up to 52 %).

At 36 °C (Fig. 3b, Fig. S8), the microbial community changed again: *Chlorobium* and *Chlorobaculum* disappeared totally. Present instead were the facultatively anaerobic, sulfur-oxidizing bacterium *Sulfuricurvum* (up to 49 %), the cyanobacterial genera *Nostoc* (up to 36 %) and *Limnothrix* (up to 25 %), and the phototrophic sulfur bacteria *Phaeospirillum* (up to 11 %).

Separate sequencing and analysis of the upper and lower layer of liquid revealed dominance of phototrophic sulfur bacteria in the lower anoxic layer and of cyanobacteria in the upper oxic layer of the columns (Fig. S9).

### Community composition (NMDS) of the water column

The effects of temperature and nutrient addition on the relative abundance described in the previous sections were very evident in the analyses of community composition via NMDS (Fig. 4a). Compositional variation along the first NMDS axis (hereafter NMDS1) was strongly associated with temperature, but not with the nutrient addition treatment, nor the interaction of the temperature and nutrient addition treatment (Fig. 4b) (Table 1). In contrast, compositional variation along NMDS2 was affected by a strong interaction between temperature and the nutrient addition treatment (Fig. 4c) (Table 1). Effects of temperature on NMDS2 scores were non-linear in both the control and the nutrient addition treatments. In the control treatment, the NMDS2 score remained constant from 12 to 24 °C and then decreased up to 36 °C. In contrast, in the nutrient addition treatment NMDS2 values first decrease (12 to 16 °C), then increase (16 to 28/32 °C) and then decrease again (32 to 36 °C).

**Fig. 4.**
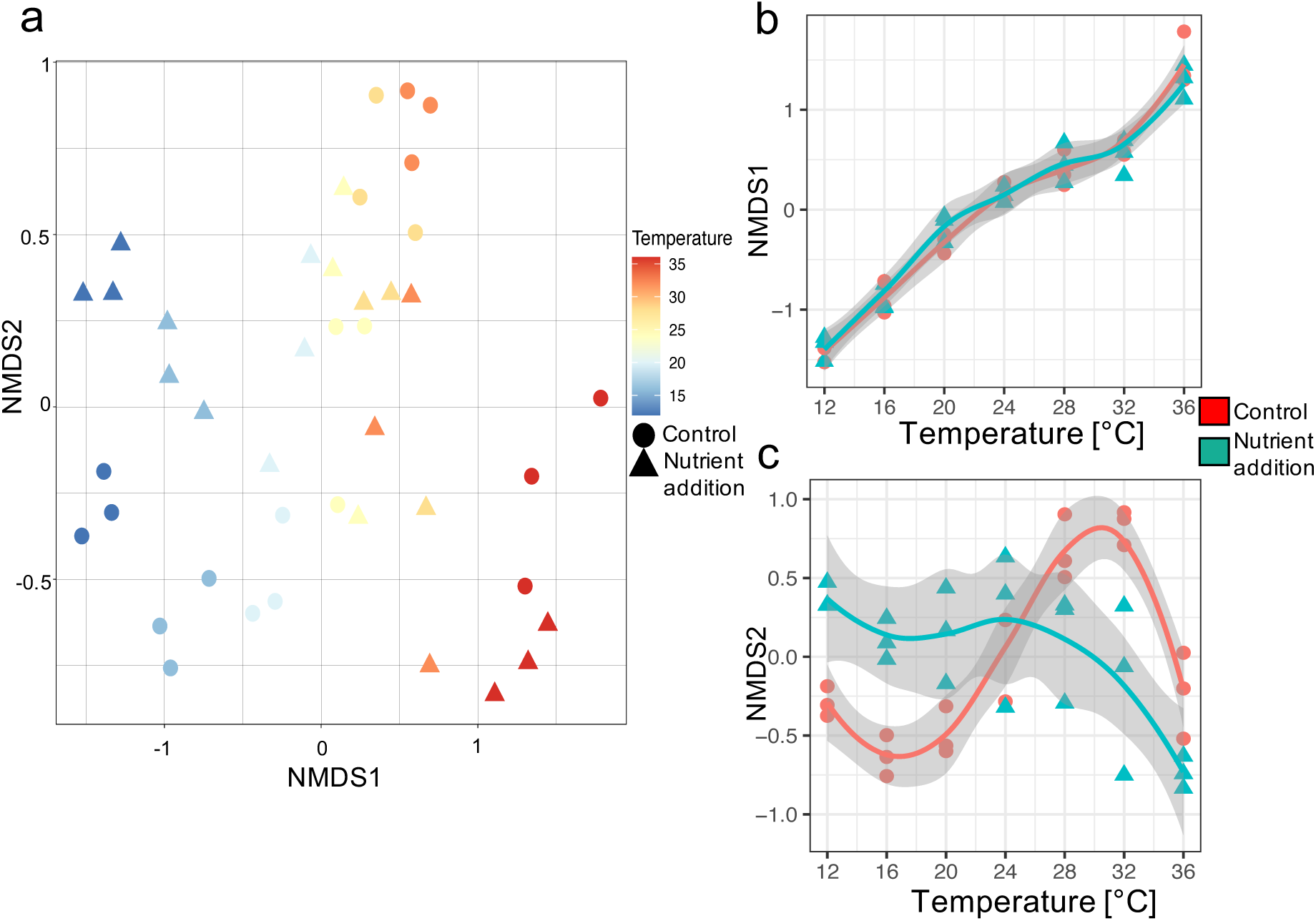
Variation in microbial community composition. (**a**) NMDS scores based on Bray-Curtis distance showing the variation in microbial community composition of the water column (**b**) The effect of temperature and nutrient addition on NMDS1. (**c**) The effect of temperature and nutrient addition on NMDS2. (**b** & **c**) Control micro-ecosystems are shown as red circles, nutrient addition-treated micro-ecosystems as blue triangles. Lines are local polynomial regressions; grey ribbons show the 95% confidence intervals.

**Fig. 5.**
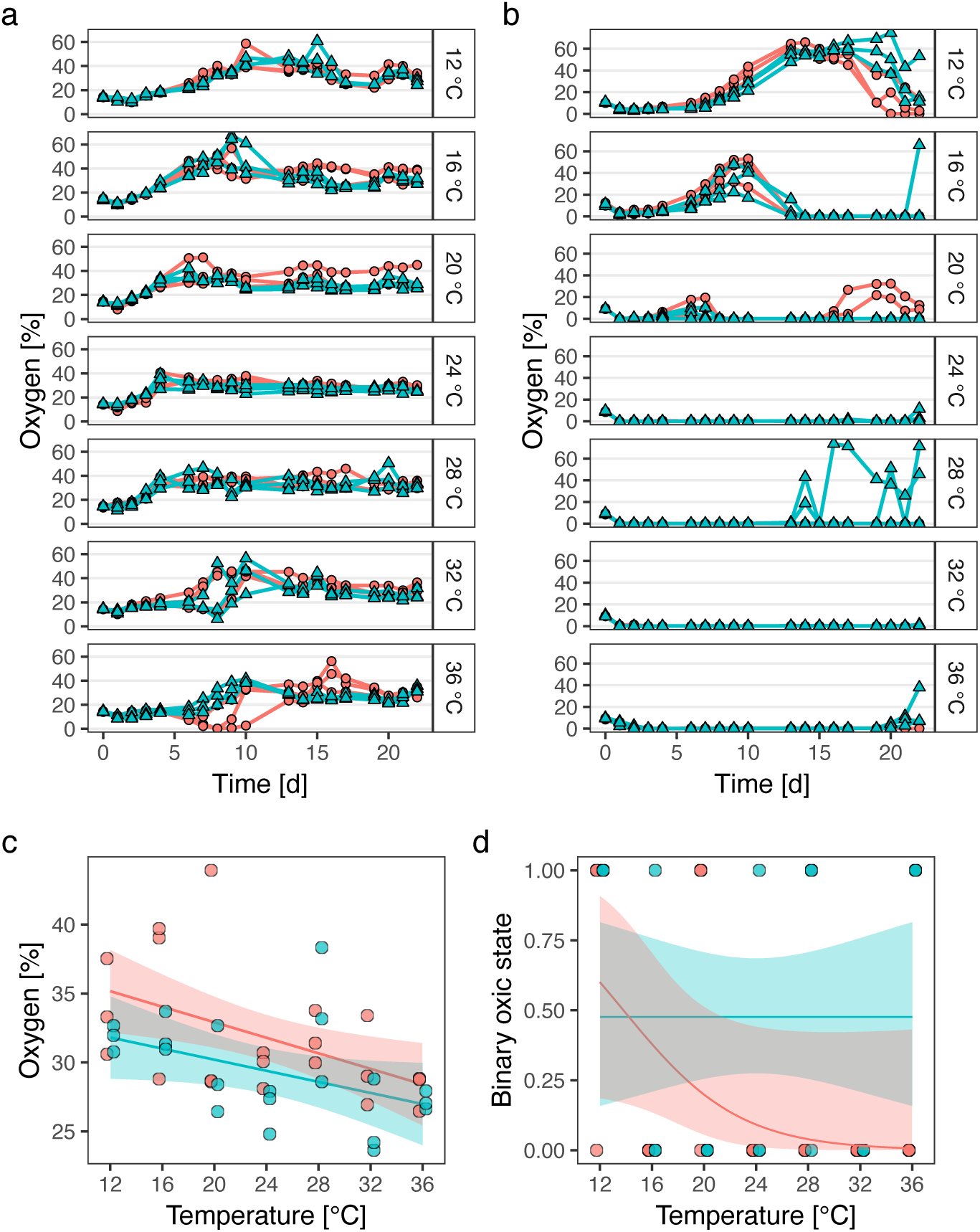
Oxygen-dynamics and long-term oxygen-states of the micro-ecosystems. In all panels, control incubations are shown in red circles (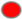), nutrient addition treatments in blue triangles (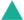). The oxygen concentration [%] of the micro-ecosystems incubated at different temperatures was measured at the top (**a**) and bottom (**b**) sensor over time. (**c**) The mean of the oxygen concentrations of the top sensor during the last three days of the experiment; lines represent the linear model fit and colored ribbons the 95 % confidence intervals. (**d**) The binary oxygen state at the bottom sensor; lines represent the regression line of the used model and colored ribbons the 95 % confidence intervals.

**Table 1.** Statistical analysis of the various response variables used in this study. Estimates of the temperature treatment are in units of the response variable per degree celcius, and estimates of the nutrient addition treatment are the effect of nutrient addition on the response variable. The interaction estimate is the effect of the nutrient addition on the estimate of the temperature effect. Terms in the main text used for p-values: no evidence p > 0.1, weak evidence 0.1 > p > 0.05, moderately strong evidence 0.05 > p > 0.01, strong evidence 0.01 > p > 0.001, very strong evidence p < 0.001. All models were general linear model with F-tests, except for the bottom oxygen analysis, which was a generalised linear model with binary response variable with likelihood ratio tests (LR-test). The total nitrogen response variable was arcsine squareroot transformed prior to analysis

|  |  | Temperature | Nutrient addition | Interaction |
| --- | --- | --- | --- | --- |
| Response variable |  |  |  |  |
| Oxygen top | Estimate | -0.28 | -4.32 | 0.08 |
|  | Standard error | 0.10 | 3.66 | 0.14 |
|  | F-test p-value | 1.9e-03 | 0.05 | 0.58 |
| Oxygen bottom | Estimate | -0.23 | -3.2 | 0.23 |
|  | Standard error | 0.12 | 2.64 | 0.13 |
|  | F-test p-value | 0.17 | 0.05 | 0.05 |
| H<sub>2</sub>S at sediment-water interphase | Estimate | -0.17 | -12.56 | 0.43 |
|  | Standard error | 8.59 | 0.25 | 0.35 |
|  | F-test p-value | 0.43 | 0.71 | 0.21 |
| pH top | Estimate | -4e-03 | 3.01 | -6e-04 |
|  | Standard error | 9.1e-03 | 0.33 | 0.01 |
|  | F-test p-value | 0.49 | <2e-16 | 0.97 |
| pH bottom | Estimate | 0.04 | -0.78 | -0.03 |
|  | Standard error | 0.01 | 0.36 | 0.01 |
|  | LR-test p-value | 2.6e-03 | 7.27e-15 | 0.07 |
| Total nitrogen (TN) | Estimate | -7.8e-04 | 0.70 | -2.7e-03 |
|  | Standard error | 1.7e-03 | 0.06 | 2.3e-03 |
|  | F-test p-value | 0.08 | < 2e-16 | 0.26 |
| Total organic carbon (TOC) | Estimate | -0.2 | 12.43 | -0.02 |
|  | Standard error | 0.45 | 15.63 | 0.63 |
|  | F-test p-value | 0.52 | 0.02 | 0.98 |
| NMDS1 | Estimate | 0.11 | 0.22 | -0.00825 |
|  | Standard error | 0.005749 | 0.21 | 0.008130 |
|  | F-test p-value | <2e-16 | 0.99 | 0.30 |
| NMDS2 | Estimate | 0.04 | 1.77 | -0.07 |
|  | Standard error | 0.01 | 0.41 | 0.02 |
|  | F-test p-value | 0.85 | 0.87 | 6.49e-05 |

### Microbial communities within the sediments

The community composition of taxonomic orders in the sediment was comparable to that in the water column (Fig. S10), though a high relative abundance of *Chlorobiales* and *Chromatiales* was detected between 20 and 32 °C. All ecosystems (except 12 °C) were rich in relative abundance of members of the anaerobic order *Spirochaetales* (up to 48 %). On genus level, however, the community consisted of a majority of sequences that could not be assigned to known taxonomic groups. With focus on sulfate-reducing microorganisms, the sediment harbored members of the order *Desulfutomaculales* up to 2 %. Besides that, genera including *Desulfobulbus*, *Desulfovibrio*, *Desulfovirga, Desulfomicrobium*, *Desulforhabdus,* were detected, but all in a low relative abundance of less than 1 %. Furthermore, members of the anaerobic biomass-degrading family *Clostridiaceae* made up to 5 % of the communities, depending on the temperature.

### Oxygen dynamics

Oxygen concentration in the top of the columns increased within the first 10 days in all incubations, and then remained oxic until the end of the experiment (Fig. 5a). Moreover, oxygen concentrations in the upper part of the columns were in general higher than 21% (concentration expected if in equilibrium with the atmosphere), and sometimes reached up to 60 %. We assume that oxygen production took place at the upper part of the columns by the algae and cyanobacteria, causing this high level of dissolved oxygen. The controls and the nutrient addition-treated columns showed only slight differences in their oxygen levels.

The long-term effect of temperature and nutrient addition on oxygen concentration in the upper part of the columns was calculated as the mean of the oxygen concentration during the last three days of the experiment (Fig. 5c). There was very strong evidence that temperature decreased the oxygen concentration from about 35 % at 12 °C to about 29 % at 36 °C (Table 1). There was moderately strong evidence that the nutrient addition treatment reduced oxygen concentration by about 4 % irrespective of temperature (i.e. there was no evidence of an interaction; Table 1).

Dynamics of the oxygen concentration of the bottom part of the micro-ecosystems were very different, and were strongly affected by temperature and nutrient addition (Fig. 5b). At 12 °C, a hump-shaped dynamic can be observed, going from anoxic to oxic and back to anoxic during the experiment. The same trend can be observed at 16°C and 20 °C, but took place earlier in the experiment. At 24-36°C there was no hump, and conditions were almost entirely anoxic, though with some rapid anoxic-oxic shifts at 16 °C, 20 °C, 24 °C, 28°C and 36 °C at the end of the experiment. These rapid shifts between oxic and anoxic states correspond with shifts in the position of the white microbial population from above (anoxic) to below (oxic) the bottom oxygen sensor (Fig. 2c).

The long-term effect of temperature and nutrient addition on the oxygen concentration of the bottom part of the columns was assessed by transforming the measured oxygen concentration into a binary variable: oxic (O_2_ concentration > 2 %) or anoxic (O_2_ concentration < 2%). There was moderate evidence of an interaction between temperature and nutrient addition in determining oxic or anoxic state (Table 1). The anoxic state became more likely as temperature increased in the control treatment, but temperature did not have such an effect in the nutrient addition treatment (Table 1).

### Chemical parameters

Highest H_2_S concentrations at the height of the bottom oxygen sensor were measured at 32 °C on day 8 (max 6 mg/L) and day 14 (max 5 mg/L), before the H_2_S concentration decreased strongly by day 20 (Fig. 6a). H_2_S on day 22 at the sediment-water interphase was detectable in most micro-ecosystems (Fig. 6b), with frequently high concentrations with more than 15 mg/L. However, there was no evidence for an effect of temperature, nutrient addition, or of their interaction on H_2_S at the end of the experiment (Table 1).

**Fig. 6.**
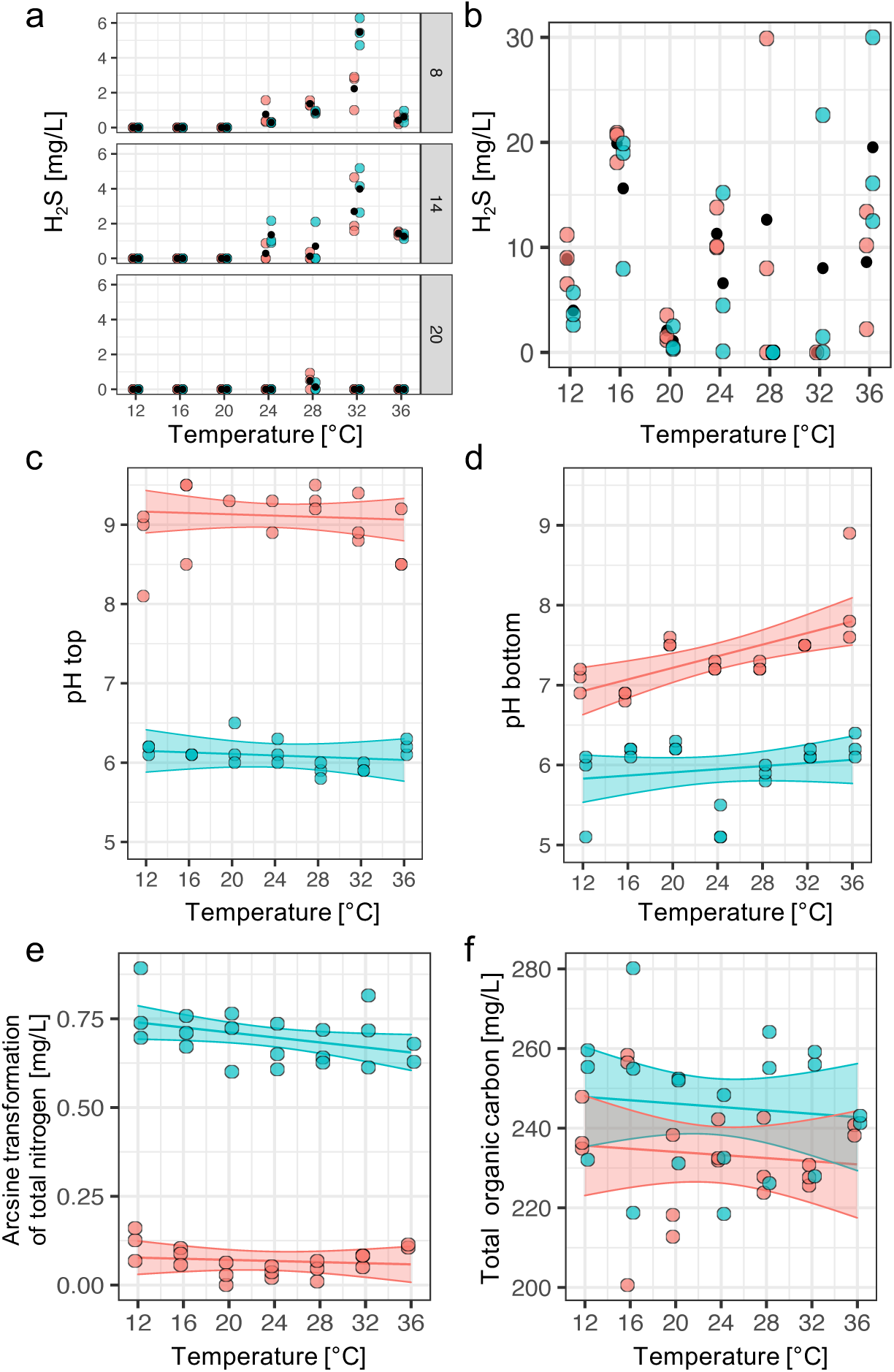
Chemical parameters of the micro-ecosystems. (**a**) Measurement of the Net-H_2_S concentration [mg/L] at the bottom oxygen sensor per temperature at incubation day 8, 14 and 20. **(b)** Measurement of the Net-H_2_S concentration [mg/L] at the soil-water interphase per temperature at final incubation day 22. (**c**) pH of the columns per temperature at the upper part. (**d**) pH of the columns per temperature at the bottom part. (**e**) Total nitrogen [mg/L] concentration [mg/L] of the micro-ecosystems depending on incubation temperature. Data were arcsine transformed for statistical analysis. (**f**) Total organic carbon concentration [mg/L] of the micro-ecosystems depending on incubation temperature. Controls are shown in red dots (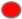), nutrient addition treatments with blue dots (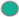). Small black points indicate mean values. Fitted models and confidence intervals (95 %) are shown as respectively lines and shaded areas for plots for which there was evidence of their significance.

There was very strong evidence of that nutrient addition affected the top (Fig. 6c) and bottom pH (Fig. 6d, Table 1), and additionally strong evidence that temperature affected the bottom pH value. Moreover, the pH of top and bottom part of the micro-ecosystems showed clear differences (compare Fig. 6c & d). While the control micro-ecosystems had a pH gradient from ∼pH 7.5 at the bottom to ∼ pH 9 at the top, no such gradients occurred in the nutrient addition-treatment, which exhibited a pH of ∼ 6 in both the top and bottom parts of the columns.

In addition, total nitrogen (TN) and total organic carbon (TOC) were analyzed for the water column of the micro-ecosystems (Fig. 6e & f), and data indicated a very strong evidence that nutrient addition increased TN concentration and a strong evidence for increasing the TOC concentration (Table 1).

## Discussion

Understanding and predicting the consequences of multifarious environmental change is very important in a broad range of research disciplines, and investigations should be made from global- to micro-scales (Balser et al. 2006). In contrast to existing work on higher-developed organisms (Tylianakis et al. 2008; González-Varo et al. 2013; Pennekamp et al. 2018) less is known about how bacterial ecosystems react to multifarious global change scenarios (Coyle et al. 2017; Hutchins and Fu 2017; Cavicchioli et al. 2019; Rillig et al. 2019). In addition, there is a need of experimental studies (in addition to natural systems and mathematical models) of complex microbial model ecosystems that can potentially display diverse responses to changing environmental conditions for various microbial groups in one experimental unit. With inspiration from Winogradsky columns (Dworkin and Gutnick 2012; Pagaling et al. 2017), we introduce a modified experimental micro-ecosystem, that includes the development of aerobic and anerobic microbial communities in one experimental ecosystem in a replicable manner. In this study we demonstrate that such an approach represents an appropriate model system for analyzing responses of complex and stratified microbial communities to global change scenarios.

Some of our results, for example the stratification and temperature effects, may be considered unsurprising and “textbook knowledge”. This is good evidence that these small and liquid dominated microbial ecosystems are appropriate analogues of naturally occurring communities. Furthermore, we used this model system to shed light on the consequences of multifarious environmental change, namely simultaneous warming and nutrient deposition—a relatively novel area of ecology, evolution, and microbiology. We hypothesized a simultaneous effect on oxygen dynamics and microbial community compositions. While temperature is considered to be a main driver for shaping soil/sediment communities (Cole et al. 2013; Garcia-Pichel et al. 2013; Deng et al. 2018), we furthermore observed non-additive effects of warming and nutrient addition on the aquatic microbial community (Fig. 4c, Table 1). This highlights the importance of studying potential interactions of multiple environmental change parameters simultaneously (Niiranen et al. 2013), given that they can differ in effect compared to the sum of individual disturbances. Environmental change cannot be reduced to the sums of individual factors. It is multifarious, providing the opportunity for “risk multiplication” and for greater challenges when attempting to understand and predict ecosystem responses to environmental change (Schulte to Bühne et al. 2020). Besides the effect on overall microbial community composition, our results additionally indicated that multiple drivers affect the oxic and anoxic layers differently. The oxic layer of the system was affected by temperature and nutrient addition additively (Fig. 5a & c, Table 1). In contrast, the anoxic layers were affected by non-additive effects of temperature and nutrient addition (Fig. 5b & c). These observations might be due to the different biogeochemistry processes in the two strata, such as the interplay of specific microbial communities, and their interplay with available nutrients, trace elements, as well as abiotic factors and the presence of specific oxygen-reducing sulfur compounds (Jorgensen et al. 1979; Chapra and Canale 1991; Luther et al. 2003; Garcia et al. 2013; Yu et al. 2014; Bush et al. 2017).

The microbial community composition and the oxygen dynamics are tightly coupled in a feedback loop. Key players in this feedback loop are oxygen-producing cyanobacteria, sulfate-reducing bacteria and phototrophic sulfur bacteria (Lee et al. 2014; Bush et al. 2017). While cyanobacteria and especially green and purple sulfur bacteria had high relative abundance in the micro-ecosystems, the relative abundance of sulfate-reducing microorganisms was comparably low. This observation could be explained by the sampling schedule: detection of high H_2_S concentration within the micro-ecosystems indicates a nutrient rich habitat for phototrophic sulfur bacteria (Guerrero et al. 1985; Hamilton et al. 2014), which suggests previous high activity of sulfate-reducing microorganisms (Sass et al. 1997; Pimenov et al. 2014). Earlier or later sampling may have revealed more oxidized sulfur-compounds and higher abundance of sulfate-reducing microorganisms. Further explanation of this unbalanced ratio between these two functional groups could probably be found in the light-dark- and the sediment-water ratio of our model system. With increasing temperature, microorganisms changed from aerobic respresentatives (*Giesbergeria*, *Uliginosibacterium*) to anaerobic ones (*Chlorobaculum*, *Chlorobium*), which is also confirmed by the oxygen measurements. Higher turnover rate of organic substrates of the microbes in the sediment and higher H_2_S production supports the dominant growth of phototrophic sulfur bacteria at moderate and higher temperatures (Nedwell et al. 1994; Findlay and Kamyshny 2017; Velthuis et al. 2018). The dominant blooms of *Chlorobium* with increasing temperature is an important finding and its ecological consequences should be investigated for future global change scenarios.

Some results hint at the presence of alternate stable states (Beisner and Cuddington 2010) in the phototrophic sulfur bacterial communities. There was considerable variation among some of the replicates of the same treatment combination, despite them having very similar initial conditions. For example, some contained either the green phototrophs *Chlorobium* or *Chlorobaculum*, or the purple phototroph *Allochromatium* (based on sequencing and macro/microscopical observation of the respective columns). Beside these alternative compositions within the same functional group, also the oxygen dynamics indicated interesting patterns, particularly at 28 °C, where two of three replicates of the nutrient addition-treated micro-ecosystems became oxic at the bottom sensor. Since no significant differences in the microbial community can be observed in these incubations, vertical movements of the oxic-anoxic interphase may have occurred, which was also visible by the different height of the white microbial community, consisting of *Rhodospirillaceae* representatives. Additionally, sinking of cyanobacteria into the anoxic layer could also result in temporary changes of oxygen dynamics in our dynamic system.

An interesting aspect could be observed regarding the pH gradient of the water column, which showed a slightly alkaline pH on the water surface in the controls, probably due to photosynthesis reactions of cyanobacteria/algae and HCO_3_^-^ in the unbuffered system, and a slightly acidic pH at the bottom, probably due to organic matter degradation. This gradient was removed by the addition of NH_4_H_2_PO_4_, which decreases the pH due to its acidic character and buffering activity on the surface to pH ∼6.

This study reveals that two environmental change factors (i.e. temperature and nutrient availability) caused large and non-additive variation in the composition of aquatic microbial communities and in the abiotic conditions of the ecosystem.The advantages of this system include high parallelization and replication, easy and non-destructive sampling, versatility of testable conditions and manipulations, and are of additional interest besides *in situ* ecosystem studies. We believe that these insights are only the tip of the iceberg of what can be learned from such model micro-ecosystems with strong and even stratified spatial environmental gradients. Even in the described study we could have included many more elements, such as characterization of the conditions at the sampling site, measurements of organic compounds composition in the water column, and a control without cellulose. Further research with this new experimental system could take many paths, including: studying the stability of the communities to press and pulse perturbations, and how this stability may depend on aspects of community composition, such as functional composition and intraspecific diversity; the extent and significance of evolutionary processes such as mutation and selection for mediating effects of environmental change; and observation of community composition via metagenomic methods, in order to capture not just the bacterial component of the micro-ecosystems, but also to research the functional significance of other likely inhabitants, such as viruses. Finally, one could research why these micro-ecosystems did not become entirely oxic or entirely anoxic, and why there was little evidence of the discontinuous responses to environmental change that are predicted for systems with strong positive feedbacks, such as this one.

## Acknowledgements

MS was funded by Forschungskredit of the UZH (FK-20-125). OLP was supported by the University of Zurich Research Priority Programs in Global Change and Biodiversity. We thank Robin Hostettler for technical support and Marcel Freund for preparing the light-gradient chambers. We thank the Functional Genomic Center Zurich for sequencing efforts and support with sequencing analysis.

The authors declare that there is no conflict of interest.

## Author contributions

OP and MS planned the experimental set-up. MS performed all experiments in the lab. MS performed the upstream bioinformatics. OP and MS performed the downstream analyses of microbial community compositions. UD supported statistical analysis and modelling of the oxygen data and general coding in RStudio. YC performed total nitrogen and organic carbon measurements and provided technical support. OP and MS drafted the manuscript. All authors confirmed the final version of the manuscript.

## References

Allison SD, Lu Y, Weihe C, et al (2013) Microbial abundance and composition influence litter decomposition response to environmental change. Ecology 94:714–725. https://doi.org/10.1890/12-1243.1

Antranikian G, Suleiman M, Schäfers C, et al (2017) Diversity of bacteria and archaea from two shallow marine hydrothermal vents from Vulcano Island. Extremophiles 21:733–742. https://doi.org/10.1007/s00792-017-0938-y

Balser TC, McMahon KD, Bart D, et al (2006) Bridging the gap between micro - and macro-scale perspectives on the role of microbial communities in global change ecology. Plant Soil 289:59–70. https://doi.org/10.1007/s11104-006-9104-5

Beisner BE, Cuddington K (2010) Alternative stable states in ecology. Front Ecol 327:259–266. https://doi.org/10.1007/s10509-010-0328-8

Boetius A (2019) Global change microbiology — big questions about small life for our future. Nat Rev Microbiol 17:331–332. https://doi.org/10.1038/s41579-019-0197-2

Bush T, Diao M, Allen RJ, et al (2017) Oxic-Anoxic regime shifts mediated by feedbacks between biogeochemical processes and microbial community dynamics. Nat Commun 8:. https://doi.org/10.1038/s41467-017-00912-x

Callahan BJ, McMurdie PJ, Rosen MJ, et al (2016) DADA2: High-resolution sample inference from Illumina amplicon data. Nat Methods 13:581–583. https://doi.org/10.1038/nmeth.3869

Cao P, Lu C, Yu Z (2018) Historical nitrogen fertilizer use in agricultural ecosystems of the contiguous United States during 1850--2015: application rate, timing, and fertilizer types. Earth Syst Sci Data 10:969–984. https://doi.org/10.5194/essd-10-969-2018

Cavicchioli R, Ripple WJ, Timmis KN, et al (2019) Scientists’ warning to humanity: microorganisms and climate change. Nat Rev Microbiol 17:569–586. https://doi.org/10.1038/s41579-019-0222-5

Chapra SC, Canale RP (1991) Long-term phenomenological model of phosphorus and oxygen for stratified lakes. Water Res 25:707–715. https://doi.org/10.1016/0043-1354(91)90046-S

Cole JK, Peacock JP, Dodsworth JA, et al (2013) Sediment microbial communities in Great Boiling Spring are controlled by temperature and distinct from water communities. ISME J 7:718–729. https://doi.org/10.1038/ismej.2012.157

Coyle DR, Nagendra UJ, Taylor MK, et al (2017) Soil fauna responses to natural disturbances, invasive species, and global climate change: Current state of the science and a call to action. Soil Biol Biochem 110:116–133. https://doi.org/10.1016/j.soilbio.2017.03.008

De Vos MGJ, Zagorski M, McNally A, Bollenbach T (2017) Interaction networks, ecological stability, and collective antibiotic tolerance in polymicrobial infections. Proc Natl Acad Sci U S A 114:10666–10671. https://doi.org/10.1073/pnas.1713372114

Deng Y, Ning D, Qin Y, et al (2018) Spatial scaling of forest soil microbial communities across a temperature gradient. Environ Microbiol 20:3504–3513. https://doi.org/10.1111/1462-2920.14303

Dworkin M (2012) Sergei Winogradsky: A founder of modern microbiology and the first microbial ecologist. FEMS Microbiol Rev 36:364–379. https://doi.org/10.1111/j.1574-6976.2011.00299.x

Dworkin M, Gutnick D (2012) Sergei Winogradsky: a founder of modern microbiology and the first microbial ecologist. FEMS Microbiol Rev 36:364–379. https://doi.org/10.1111/j.1574-6976.2011.00299.x

Evans SE, Wallenstein MD (2014) Climate change alters ecological strategies of soil bacteria. Ecol Lett 17:155–164. https://doi.org/10.1111/ele.12206

Falkowski PG, Fenchel T, Delong EF (2008) The microbial engines that drive earth’s biogeochemical cycles. Science (80-) 320:1034–1039. https://doi.org/10.1126/science.1153213

Findlay AJ, Kamyshny A (2017) Turnover Rates of Intermediate Sulfur Species (Sx2-, S0, S2O32-, S4O62-, SO32-) in Anoxic Freshwater and Sediments. Front Microbiol 8:2551. https://doi.org/10.3389/fmicb.2017.02551

Garcia-Pichel F, Loza V, Marusenko Y, et al (2013) Temperature Drives the Continental-Scale Distribution of Key Microbes in Topsoil Communities. Science (80-) 340:1574–1577. https://doi.org/10.1126/science.1236404

Garcia SL, Salka I, Grossart H-P, Warnecke F (2013) Depth-discrete profiles of bacterial communities reveal pronounced spatio-temporal dynamics related to lake stratification. Environ Microbiol Rep 5:549–555. https://doi.org/10.1111/1758-2229.12044

Gloor GB, Macklaim JM, Pawlowsky-Glahn V, Egozcue JJ (2017) Microbiome Datasets Are Compositional: And This Is Not Optional. Front Microbiol 8:2224. https://doi.org/10.3389/fmicb.2017.02224

González-Varo JP, Biesmeijer JC, Bommarco R, et al (2013) Combined effects of global change pressures on animal-mediated pollination. Trends Ecol Evol 28:524–530. https://doi.org/10.1016/j.tree.2013.05.008

Guerrero R, Montesinos E, Pedrós-Alió C, et al (1985) Phototrophic sulfur bacteria in two Spanish lakes: Vertical distribution and limiting factors1. Limnol Oceanogr 30:919–931. https://doi.org/10.4319/lo.1985.30.5.0919

Hamilton TL, Bovee RJ, Thiel V, et al (2014) Coupled reductive and oxidative sulfur cycling in the phototrophic plate of a meromictic lake. Geobiology 12:451–468. https://doi.org/10.1111/gbi.12092

Hutchins DA, Fu F (2017) Microorganisms and ocean global change. Nat Microbiol 2:. https://doi.org/10.1038/nmicrobiol.2017.58

Hutchins DA, Jansson JK, Remais J V, et al (2019) Climate change microbiology — problems and perspectives. Nat Rev Microbiol 17:391–396. https://doi.org/10.1038/s41579-019-0178-5

Jorgensen BB, Kuenen JG, Cohen Y (1979) Microbial transformations of sulfur compounds in a stratified lake (Solar Lake, Sinai)1. Limnol Oceanogr 24:799–822. https://doi.org/10.4319/lo.1979.24.5.0799

Kertesz MA (2000) Riding the sulfur cycle - Metabolism of sulfonates and sulfate esters in Gram-negative bacteria. FEMS Microbiol Rev 24:135–175. https://doi.org/10.1016/S0168-6445(99)00033-9

Lahti L, Salojärvi J, Salonen A, et al (2014) Tipping elements in the human intestinal ecosystem. Nat Commun 5:4344. https://doi.org/10.1038/ncomms5344

Lee JZ, Burow LC, Woebken D, et al (2014) Fermentation couples Chloroflexi and sulfate-reducing bacteria to Cyanobacteria in hypersaline microbial mats. Front Microbiol 5:1–17. https://doi.org/10.3389/fmicb.2014.00061

Luek A, Rowan DJ, Rasmussen JB (2017) N-P Fertilization Stimulates Anaerobic Selenium Reduction in an End-Pit Lake. Sci Rep 7:10502. https://doi.org/10.1038/s41598-017-11095-2

Luther GW, Glazer B, Ma S, et al (2003) Iron and sulfur chemistry in a stratified lake: Evidence for iron-rich sulfide complexes. Aquat Geochemistry 9:87–110. https://doi.org/10.1023/B:AQUA.0000019466.62564.94

McMurdie PJ, Holmes S (2013) Phyloseq: An R Package for Reproducible Interactive Analysis and Graphics of Microbiome Census Data. PLoS One 8:. https://doi.org/10.1371/journal.pone.0061217

Morrison JM, Baker KD, Zamor RM, et al (2017) Spatiotemporal analysis of microbial community dynamics during seasonal stratification events in a freshwater lake (Grand Lake, OK, USA). PLoS One 12:1–29. https://doi.org/10.1371/journal.pone.0177488

Nedwell DB, Blackburn TH, Wiebe WJ (1994) Dynamic nature of the turnover of organic carbon, nitrogen and sulphur in the sediments of a Jamaican mangrove forest. Mar Ecol Prog Ser 110:223–231

Niiranen S, Yletyinen J, Tomczak MT, et al (2013) Combined effects of global climate change and regional ecosystem drivers on an exploited marine food web. Glob Chang Biol 19:3327–3342. https://doi.org/10.1111/gcb.12309

Oksanen J, Blanchet FG, Friendly M, et al (2019) Package “vegan” Title Community Ecology Package. Community Ecol Packag 2:1–297

Overmann J, van Gemerden H (2000) Microbial interactions involving sulfur bacteria: implications for the ecology and evolution of bacterial communities. FEMS Microbiol Rev 24:591–599. https://doi.org/10.1016/S0168-6445(00)00047-4

Pagaling E, Vassileva K, Mills CG, et al (2017) Assembly of microbial communities in replicate nutrient-cycling model ecosystems follows divergent trajectories, leading to alternate stable states. Environ Microbiol 19:3374–3386. https://doi.org/10.1111/1462-2920.13849

Pennekamp F, Pontarp M, Tabi A, et al (2018) Biodiversity increases and decreases ecosystem stability. Nature 563:109–112. https://doi.org/10.1038/s41586-018-0627-8

Pimenov NV, Zakharova EE, Bryukhanov AL, et al (2014) Activity and structure of the sulfate-reducing bacterial community in the sediments of the southern part of Lake Baikal. Microbiology 83:47–55. https://doi.org/10.1134/S0026261714020167

Posch T, Köster O, Salcher MM, Pernthaler J (2012) Harmful filamentous cyanobacteria favoured by reduced water turnover with lake warming. Nat Clim Chang 2:809–813. https://doi.org/10.1038/nclimate1581

Quast C, Pruesse E, Yilmaz P, et al (2012) The SILVA ribosomal RNA gene database project: improved data processing and web-based tools. Nucleic Acids Res 41:D590–D596. https://doi.org/10.1093/nar/gks1219

Rillig MC, Ryo M, Lehmann A, et al (2019) The role of multiple global change factors in driving soil functions and microbial biodiversity. Science (80-) 366:886–890. https://doi.org/10.1126/science.aay2832

Rundell EA, Banta LM, Ward D V., et al (2014) 16S rRNA Gene Survey of Microbial Communities in winogradsky columns. PLoS One 9:. https://doi.org/10.1371/journal.pone.0104134

Sass H, Cypionka H, Babenzien H-D (1997) Vertical distribution of sulfate-reducing bacteria at the oxic-anoxic interface in sediments of the oligotrophic Lake Stechlin. FEMS Microbiol Ecol 22:245–255. https://doi.org/10.1111/j.1574-6941.1997.tb00377.x

Savvichev AS, Babenko V V, Lunina ON, et al (2018) Sharp water column stratification with an extremely dense microbial population in a small meromictic lake, Trekhtzvetnoe. Environ Microbiol 20:3784–3797. https://doi.org/10.1111/1462-2920.14384

Schulte to Bühne H, Tobias JA, Durant SM, Pettorelli N (2020) Improving Predictions of Climate Change–Land Use Change Interactions. Trends Ecol Evol. https://doi.org/10.1016/j.tree.2020.08.019

Stein LY, Klotz MG (2016) The nitrogen cycle. Curr Biol 26:R94–R98. https://doi.org/10.1016/j.cub.2015.12.021

Suleiman M, Klippel B, Busch P, et al (2019) Enrichment of anaerobic heterotrophic thermophiles from four Azorean hot springs revealed different community composition and genera abundances using recalcitrant substrates. Extremophiles 23:277–281. https://doi.org/10.1007/s00792-019-01079-7

Tylianakis JM, Didham RK, Bascompte J, Wardle DA (2008) Global change and species interactions in terrestrial ecosystems. Ecol Lett 11:1351–1363. https://doi.org/10.1111/j.1461-0248.2008.01250.x

Velthuis M, Kosten S, Aben R, et al (2018) Warming enhances sedimentation and decomposition of organic carbon in shallow macrophyte-dominated systems with zero net effect on carbon burial. Glob Chang Biol 24:5231–5242. https://doi.org/10.1111/gcb.14387

Vigneron A, Cruaud P, Culley AI, et al (2021) Genomic evidence for sulfur intermediates as new biogeochemical hubs in a model aquatic microbial ecosystem. Microbiome 9:46. https://doi.org/10.1186/s40168-021-00999-x

Weiss JV, Emerson D, Backer SM, Megonigal JP (2003) Enumeration of Fe(II)-oxidizing and Fe(III)-reducing bacteria in the root zone of wetland plants: Implications for a rhizosphere iron cycle. Biogeochemistry 64:77–96. https://doi.org/10.1023/A:1024953027726

Widder S, Allen RJ, Pfeiffer T, et al (2016) Challenges in microbial ecology: Building predictive understanding of community function and dynamics. ISME J 10:2557–2568. https://doi.org/10.1038/ismej.2016.45

Yu Z, Yang J, Amalfitano S, et al (2014) Effects of water stratification and mixing on microbial community structure in a subtropical deep reservoir. Sci Rep 4:1–7. https://doi.org/10.1038/srep05821

Zavarzin GA (2006) Winogradsky and modern microbiology. Microbiology 75:501– 511. https://doi.org/10.1134/S0026261706050018

