## Supplementary Information for "Large and interacting effects of temperature and nutrient addition on stratified microbial ecosystems in a small, replicated, and liquid dominated Winogradsky column approach"

**Large and interacting effects of temperature and nutrient addition on stratified microbial ecosystems — results from an aquatic model micro-ecosystem**

Marcel Suleiman, Yves Choffat, Uriah Daugaard, and Owen Petchey

Department of Evolutionary Biology and Environmental Studies, University of Zurich, Switzerland

Corresponding author: Marcel Suleiman, Department of Evolutionary Biology and Environmental Studies, University of Zurich, Switzerland

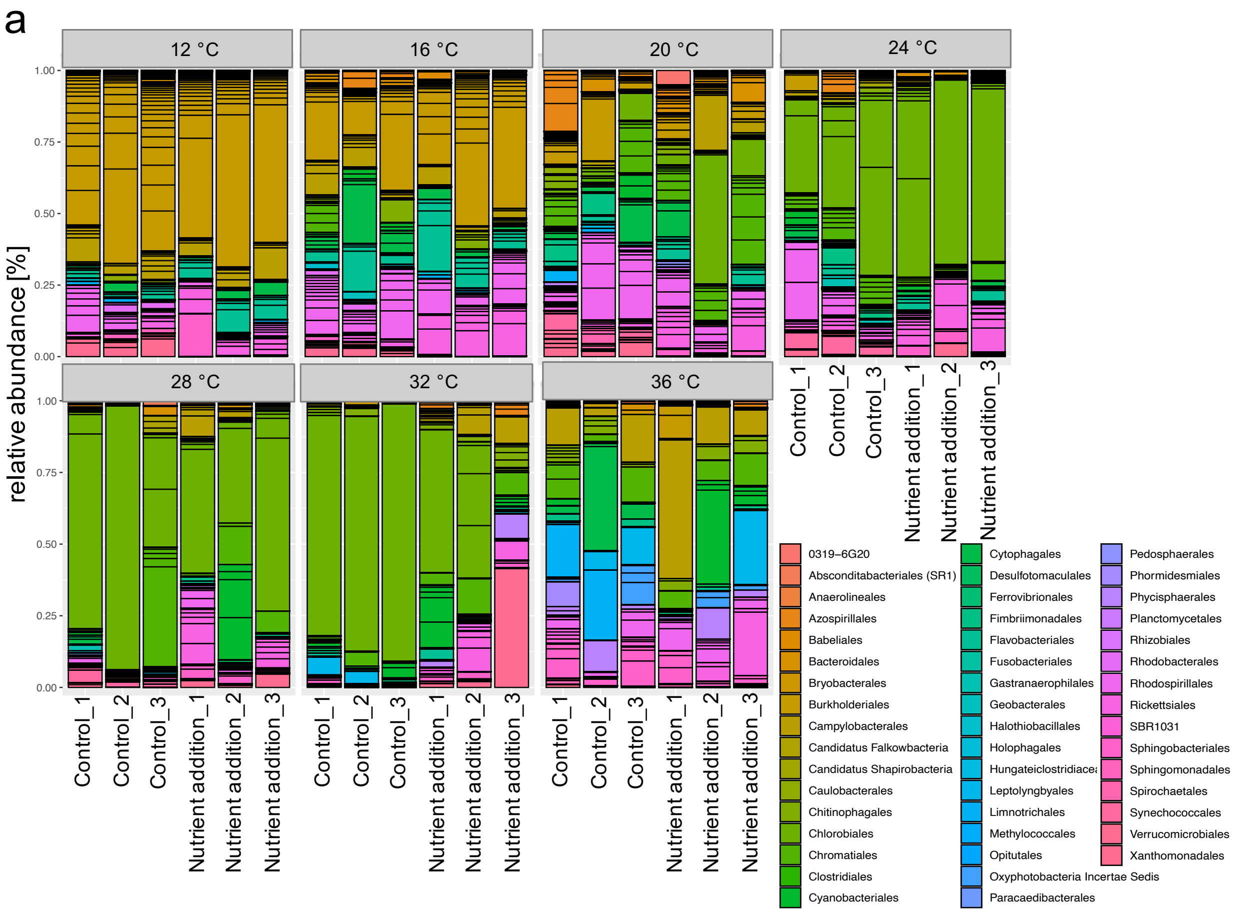


**Fig. S1 Microbial community composition of the liquid part of all micro-ecosystems.** Relative abundance on order level is shown for each micro-ecosystem.

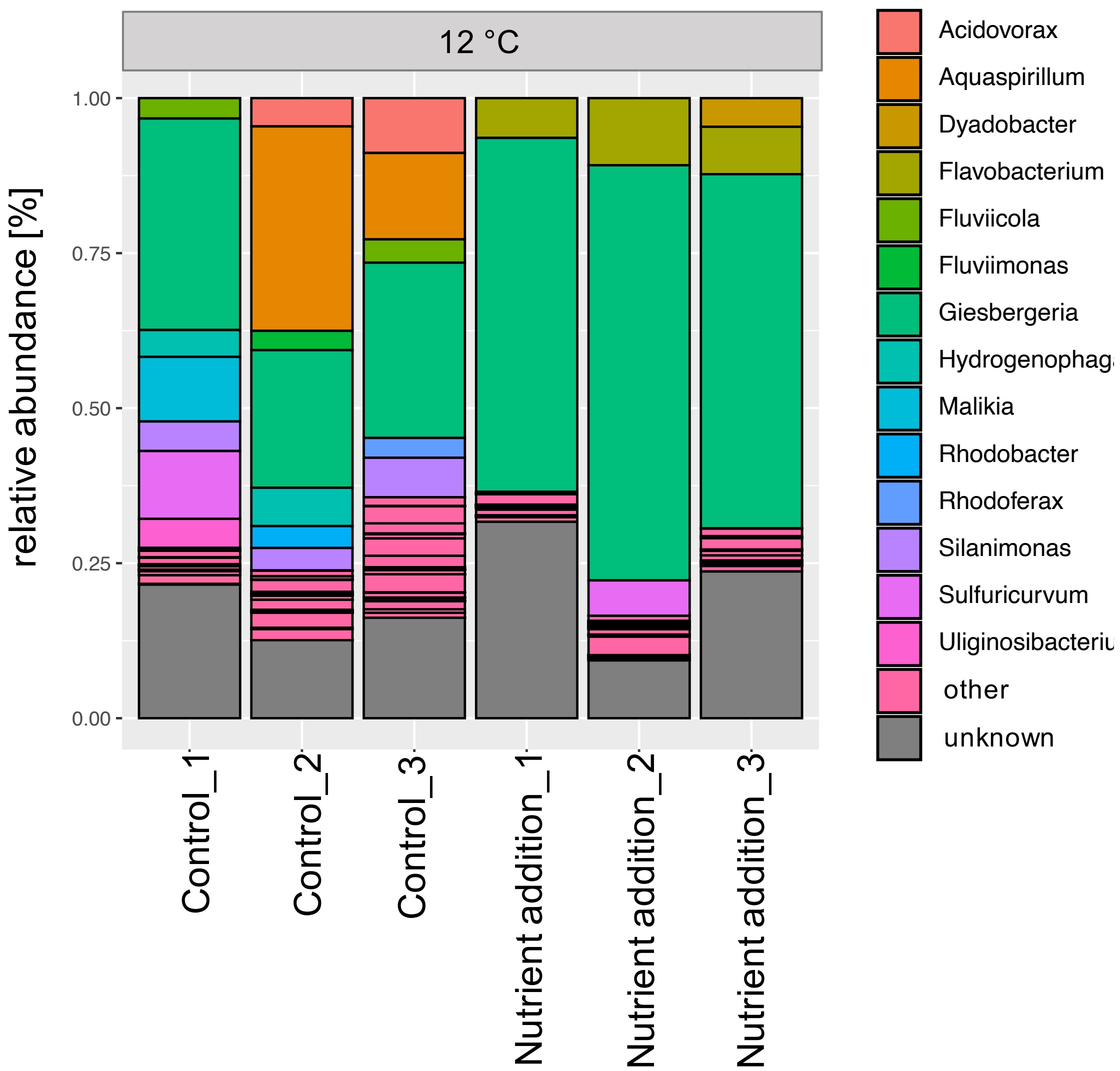


**Fig. S2 Microbial community composition of the liquid part of the micro-ecosystems incubated at 12 °C.** Relative abundance on genus level is shown for each micro-ecosystem. Relative abundances of specific genera < 3 % were assigned to “other”.

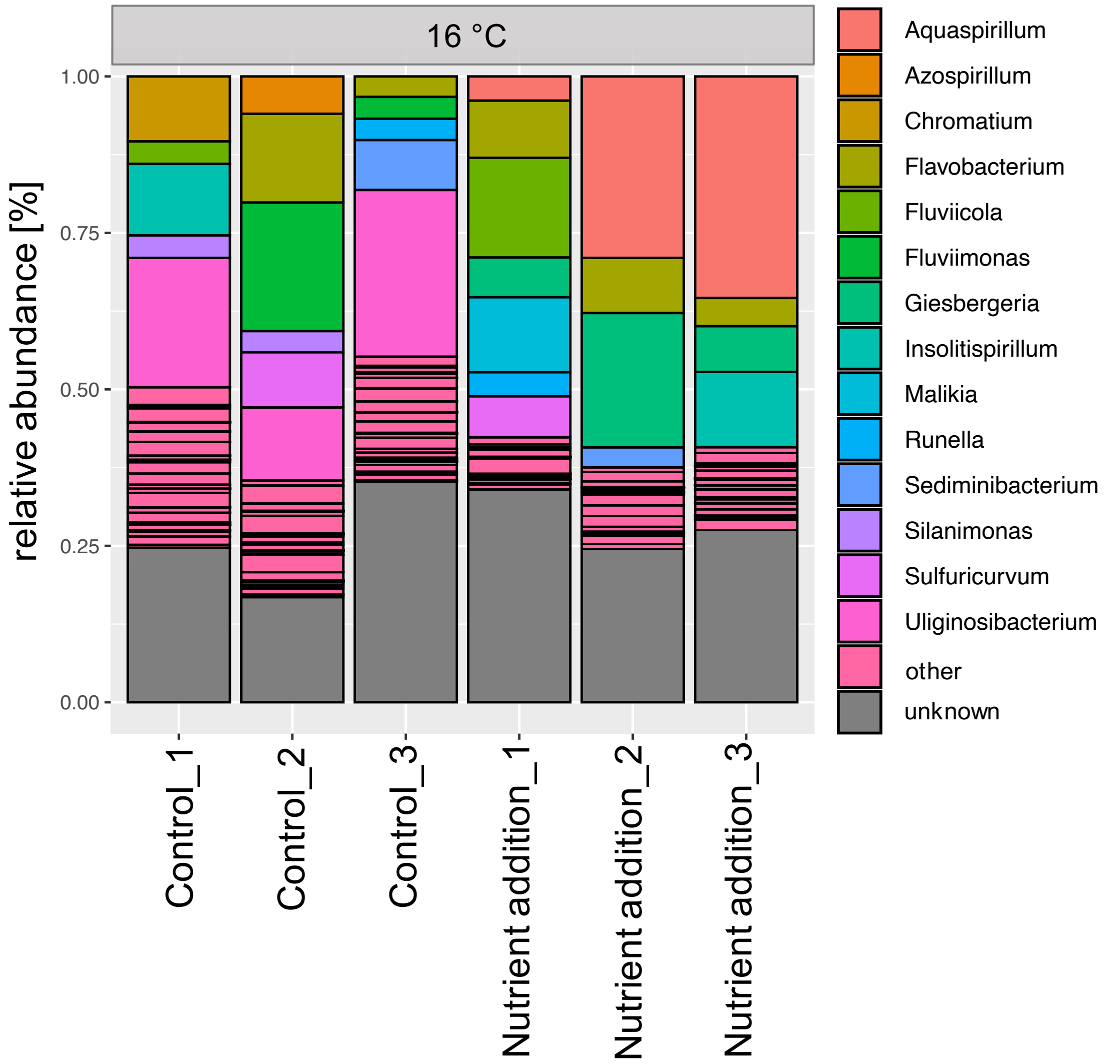


**Fig. S3 Microbial community composition of liquid part of the micro-ecosystems incubated at 16 °C.** Relative abundance on genus level is shown for each micro-ecosystem. Relative abundances of specific genera < 3 % were assigned to “other”.

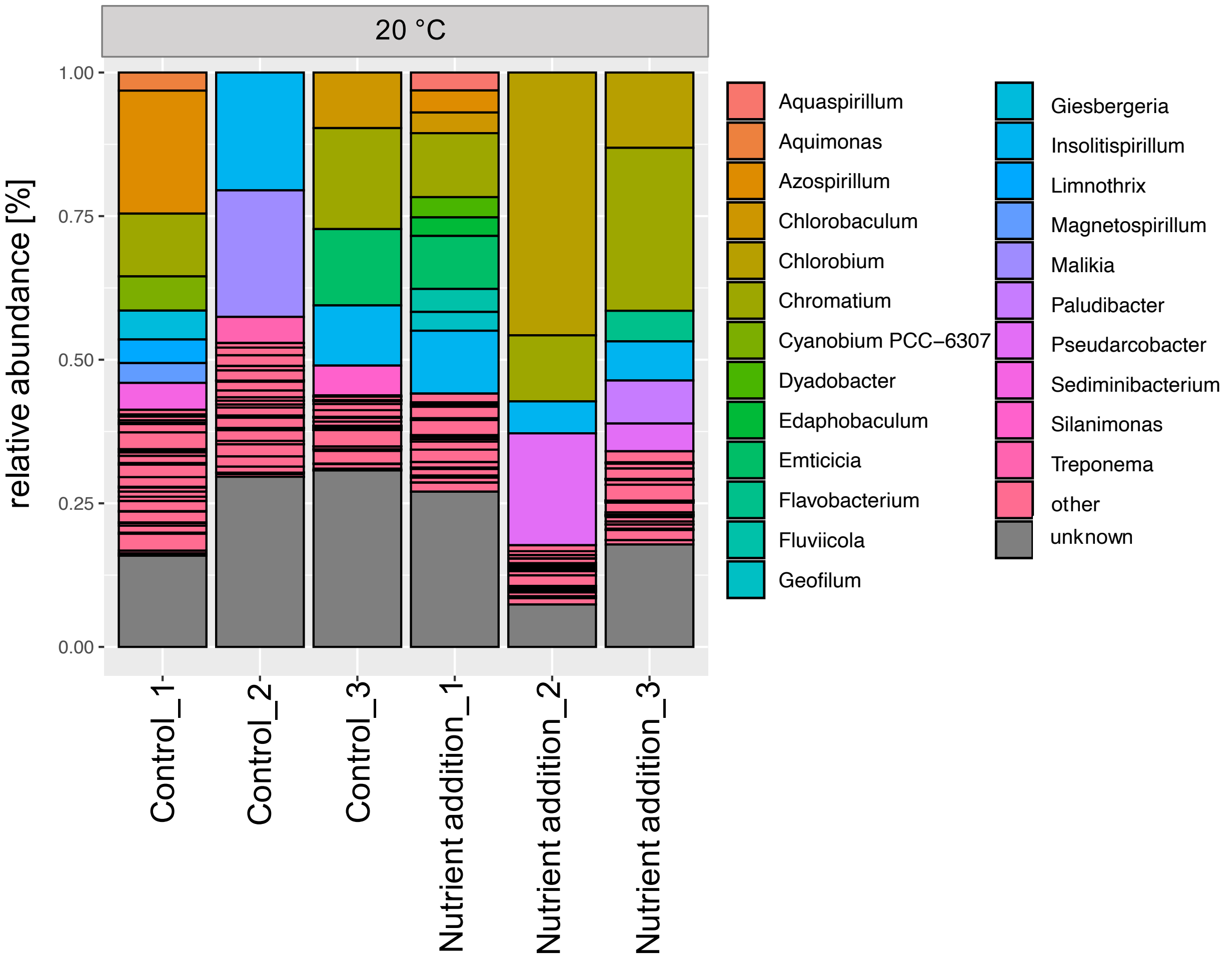


**Fig. S4 Microbial community composition of liquid part of the micro-ecosystems incubated at 20 °C.** Relative abundance on genus level is shown for each micro-ecosystem. Relative abundances of specific genera < 3 % were assigned to “other”.

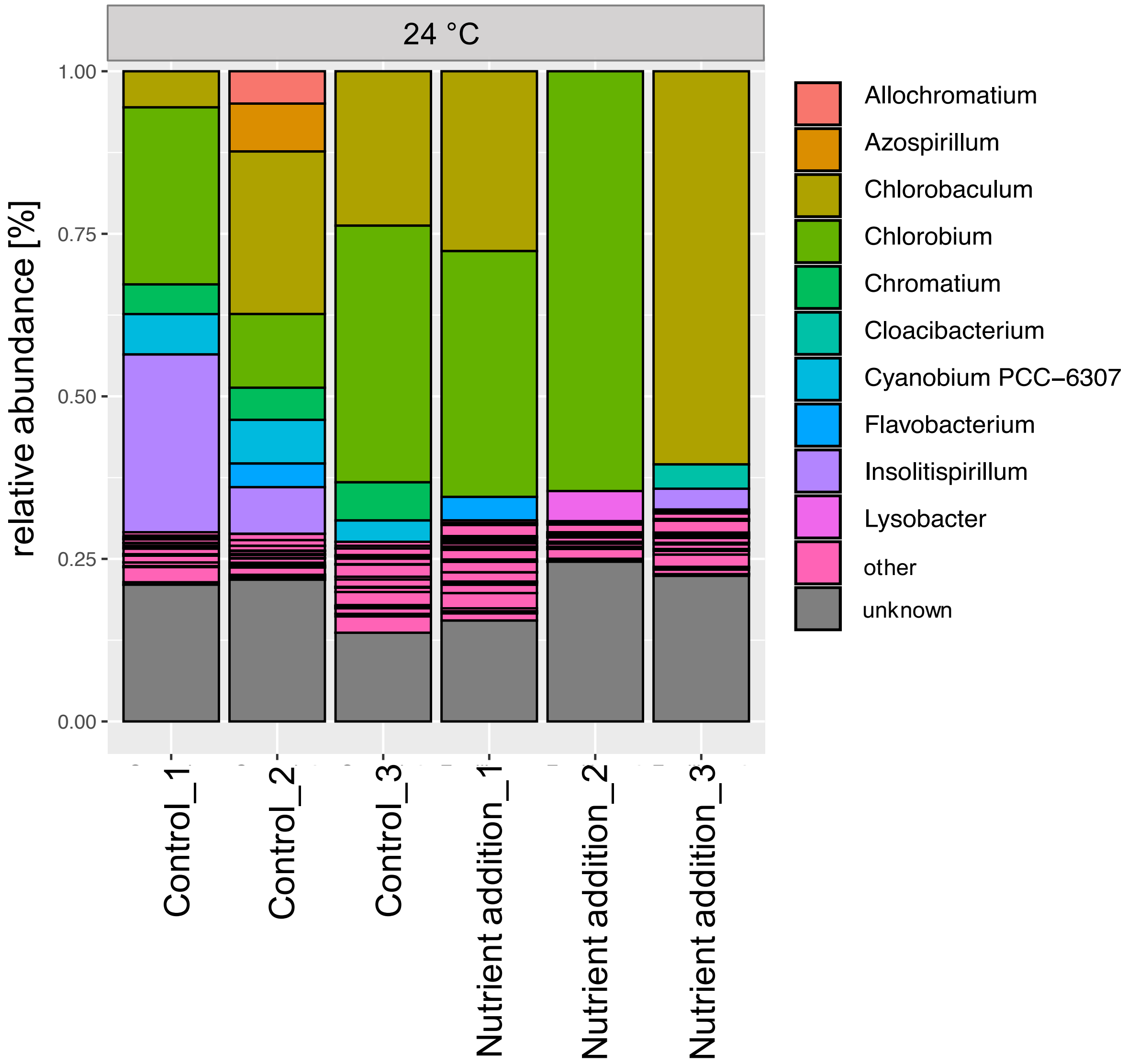


**Fig. S5 Microbial community composition of liquid part of the micro-ecosystems incubated at 24 °C.** Relative abundance on genus level is shown for each micro-ecosystem. Relative abundances of specific genera < 3 % were assigned to “other”.

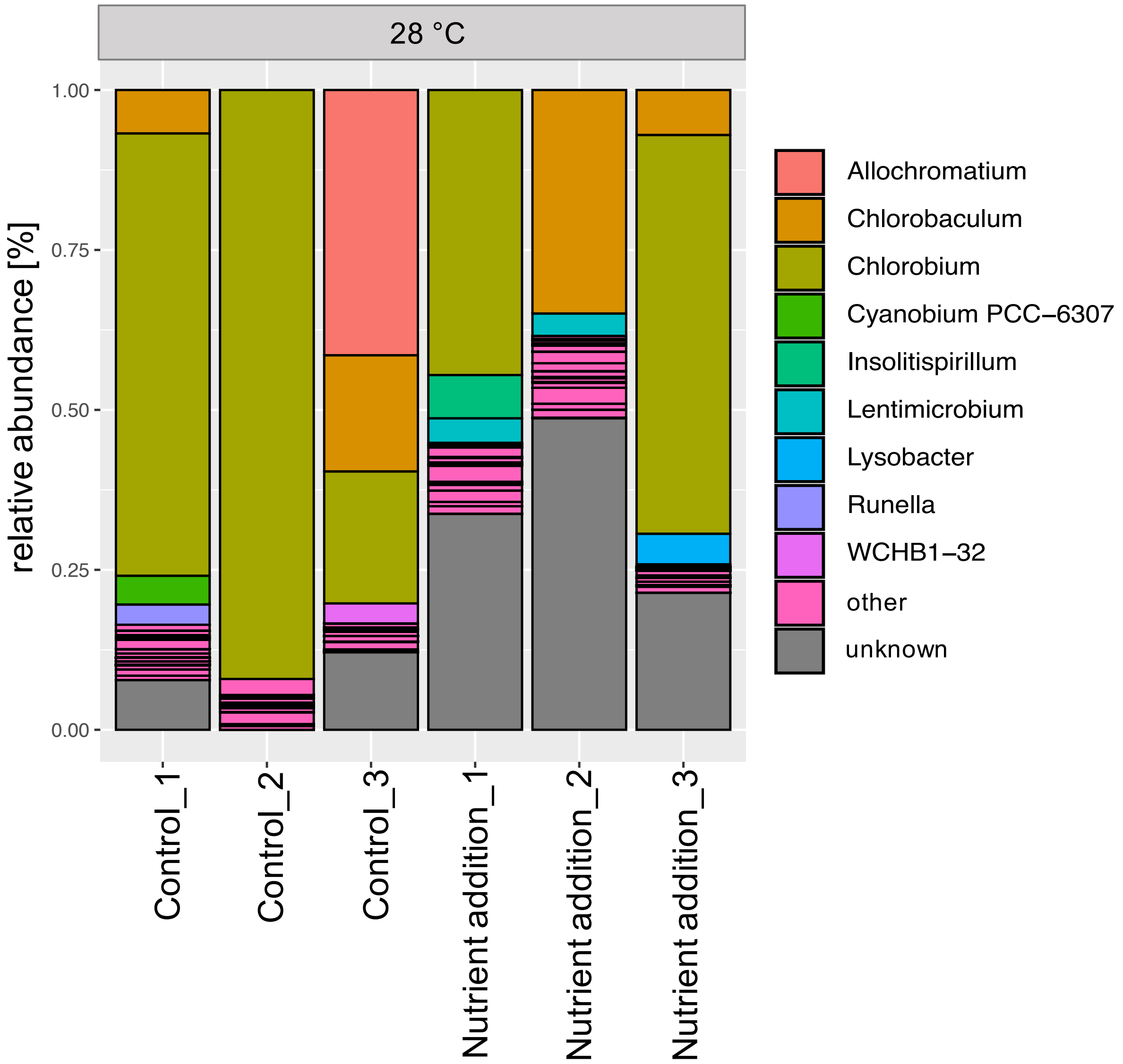


**Fig. S6 Microbial community composition of liquid part of the micro-ecosystems incubated at 28 °C.** Relative abundance on genus level is shown for each micro-ecosystem. Relative abundances of specific genera < 3 % were assigned to “other”.

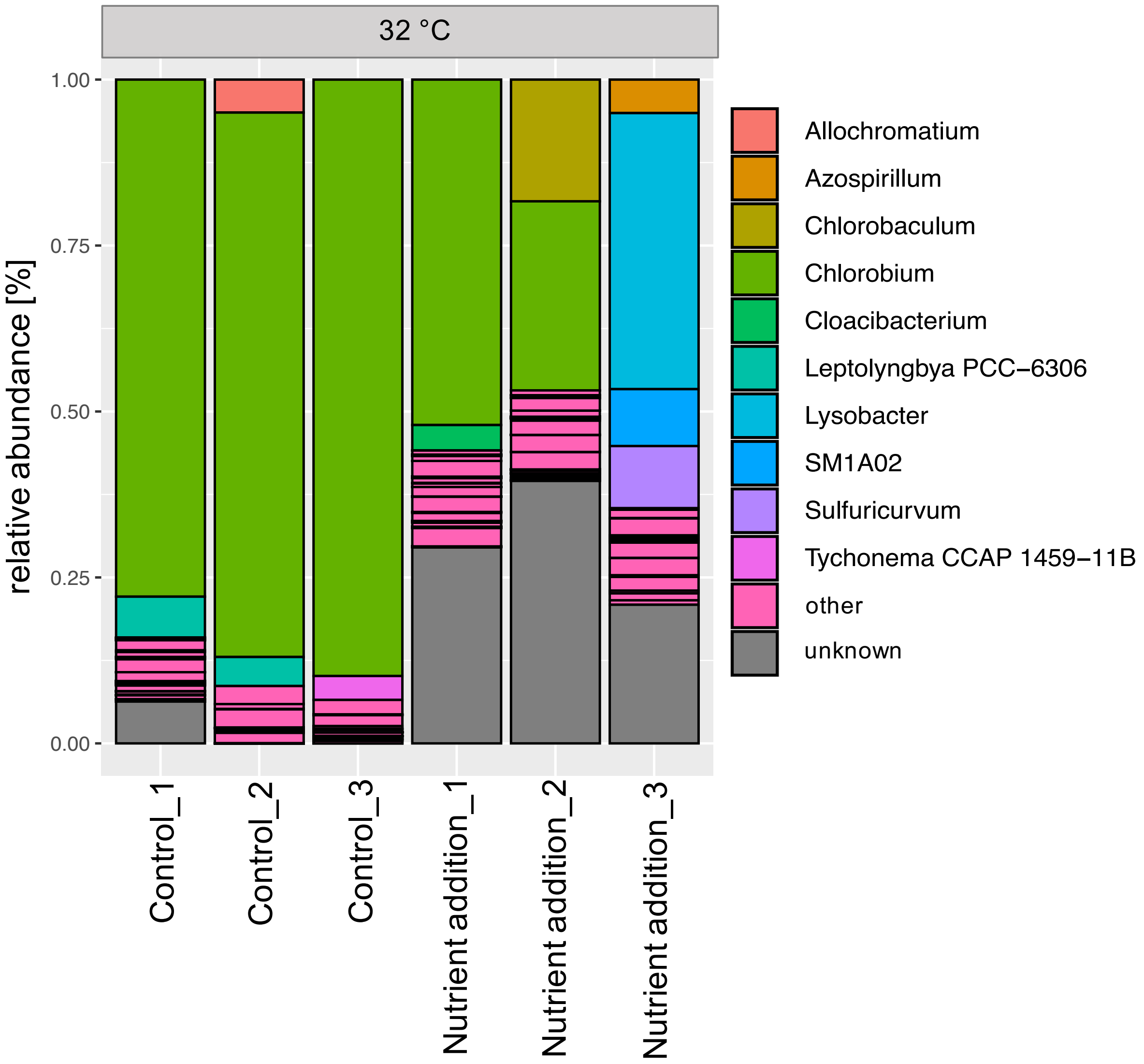


**Fig. S7 Microbial community composition of liquid part of the micro-ecosystems incubated at 32 °C.** Relative abundance on genus level is shown for each micro-ecosystem. Relative abundances of specific genera < 3 % were assigned to “other”.

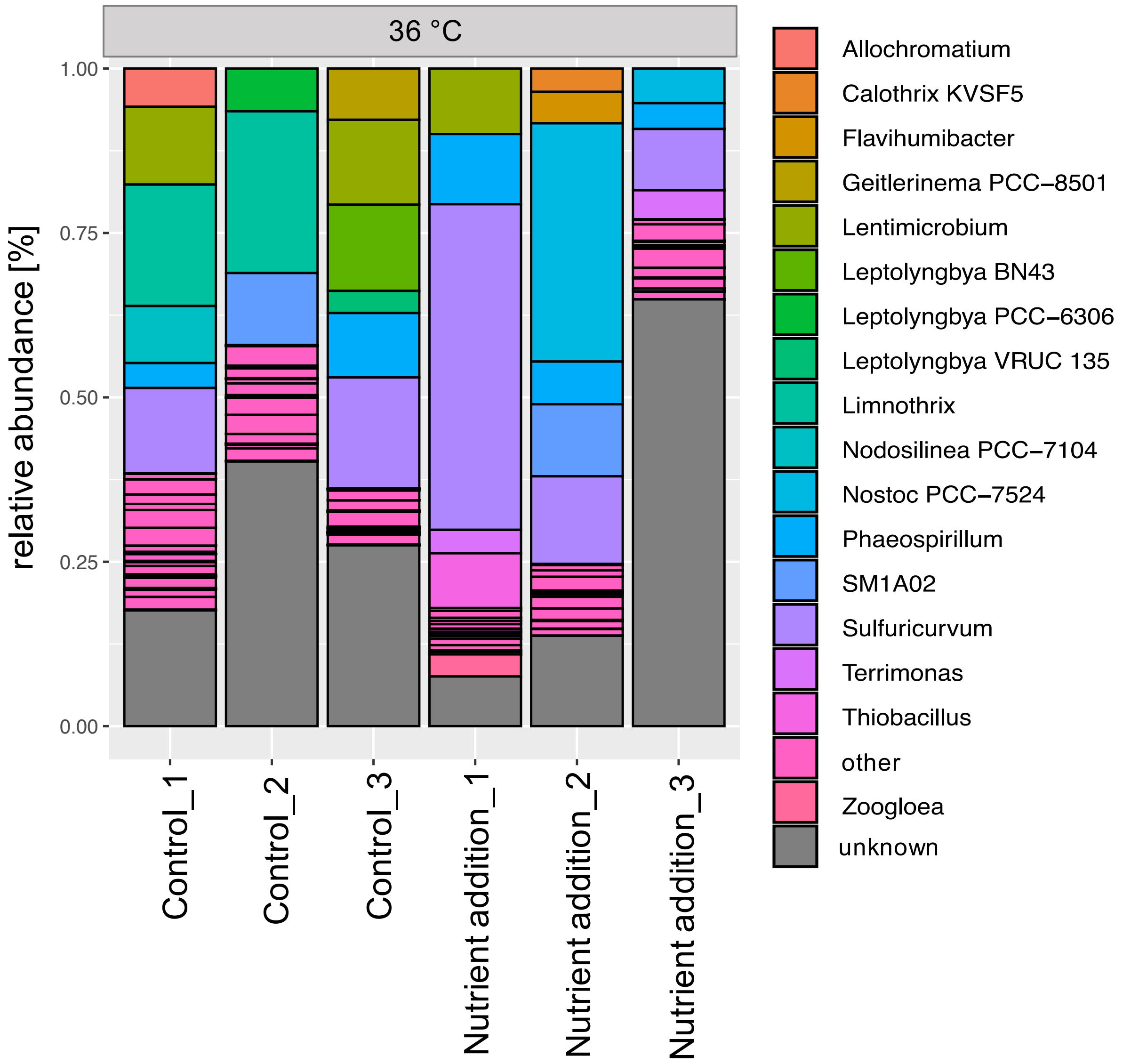


**Fig. S8 Microbial community composition of liquid part of the micro-ecosystems incubated at 36°C.** Relative abundance on genus level is shown for each micro-ecosystem. Relative abundances of specific genera < 3 % were assigned to “other”.

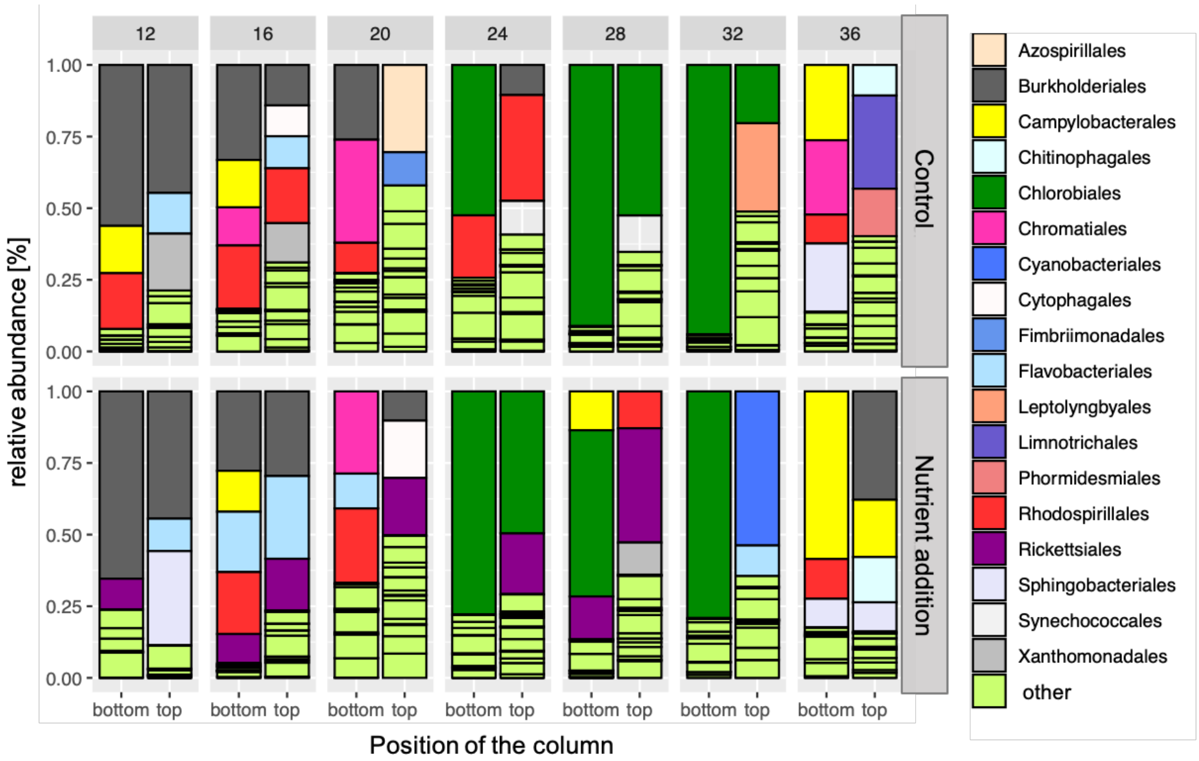


**Fig. S9 Microbial community composition of the upper and lower liquid part of replicate 1 of all treatments.** The liquid part was separated at height of the bottom oxygen layer and both layers were sequenced separately. Relative abundances of specific orders < 10 % were assigned to “other”.

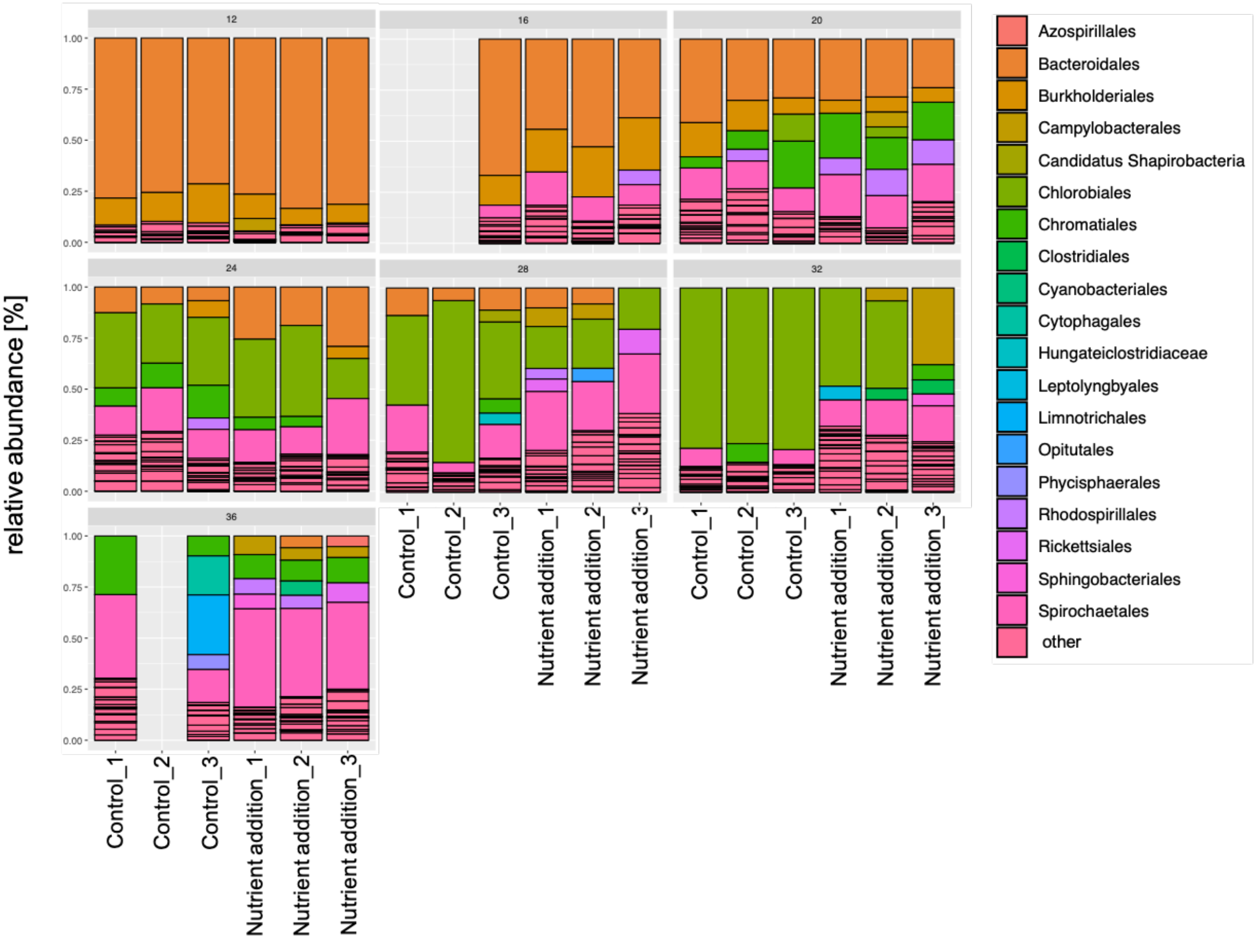


**Fig. S10 Microbial community composition of the sediments of the micro-ecosystems.** Relative abundance on order level is shown for each micro-ecosystem. Three sequencing reactions were not successful. Relative abundances of specific order < 10 % were assigned to “other”.
